# Repetitive DNA-resolved epigenomics maps chromatin rewiring during aging and malignant transformation

**DOI:** 10.64898/2026.04.29.721644

**Authors:** Juan José Alba-Linares, Juan Ramón Tejedor, Lidia Sainz-Ledo, Agustín F. Fernández, Raúl F. Pérez, Mario F. Fraga

## Abstract

Repetitive DNA constitutes more than half of the human genome, yet its chromatin landscape remains poorly understood. Here, we integrated multi-mapping reads across 3, 744 repeat subfamilies and 937 ChIP-seq experiments for six canonical histone marks in hematopoietic cells and chronic malignancies from the BLUEPRINT consortium. Chromatin signals at repeats encode cell type-specific signatures, with certain elements (Alu, SVA, ERV/Gypsy LTRs) exhibiting conserved epigenomic profiles distinct from surrounding single-copy sequences, and LTRs acting as chromatin boundaries. Combinatorial analyses further revealed five major repeat epitypes (promoter-like, active/poised enhancers, and two heterochromatin domains) that enhance annotations from uniquely-mapping reads. Beyond constitutive heterochromatin, we uncovered a highly plastic heterochromatin characterized by concurrent H3K36me3 and H3K9me3 that can dynamically switch to active epitypes in B-cell chronic neoplasms. Although malignant transformation involved global loss of histone marks across repeats, we identified targeted heterochromatinization of satellites and LTRs, mirroring changes in aged B cells, and gains of active marks at CD34⁺ enhancers and disease-related genes. Likewise, aging primes the heterochromatin landscape toward malignancy but still requires *de novo* acquisition of active marks at regulatory regions to promote transformation. Our atlas provides the most comprehensive resource for dissecting chromatin dynamics at repetitive DNA in hematopoietic aging and cancer.

## 1. INTRODUCTION

Complex eukaryotic genomes comprise both repetitive and single-copy DNA sequences, with repetitive elements accounting for more than half of the human genome (1). Once dismissed as “junk” DNA (2), these elements are now recognized as major drivers of genomic variation (3, 4) and powerful forces shaping species evolution (5–7) and disease (8). Beyond these roles, repetitive DNA actively contributes to transcriptional programs during mammalian development, particularly in embryonic totipotent states (9–11), mediates trans-generational epigenetic inheritance (12, 13), and underpins tissue-specific regulatory networks (14, 15), especially in the hematopoietic lineage (16), which can become co-opted or dysregulated in cancer (17).

This functional diversity stems from the extensive heterogeneity of repetitive elements, which are classified by genomic organization and sequence features (18, 19). Tandem repeats, including simple repeats and satellites, form head-to-tail sequence arrays typically enriched in (sub-)telomeric and (peri-)centromeric regions of eukaryotic genomes. In contrast, interspersed repeats propagate via insertion mechanisms into the host genome, and include RNA-derived pseudogenes (i.e. rRNA, tRNA, snRNA, scRNA) and transposable elements (TEs). Based on their mobilization mechanisms, TEs are further divided into class I elements, which transpose via an RNA intermediate (retrotransposons), and class II elements, which transpose as DNA (e.g., hAT, TcMar, and Helitron families). Retrotransposons are further subdivided into long terminal repeat (LTR) elements and non-LTR elements, the latter comprising long interspersed elements (LINEs), short interspersed elements (SINEs), and SINE–VNTR–Alu (SVA) retroposon elements. Among these, only full-length LINE-1 (L1) and intact LTR elements retain autonomous mobility in the human genome, whereas SINEs and SVAs rely on the L1-encoded enzymatic machinery. In general, repetitive elements and their variant copies are grouped into subfamilies based on consensus sequences, which in turn associate with larger families (e.g., L1) and broader categories (e.g., LINEs) by similarity. This complexity underscores the need for computational approaches specifically optimized for the analysis of repetitive DNA (19), including methods capable of handling multimapping reads, which can uncover patterns that have remained undetected when relying solely on uniquely mapping reads (16).

Despite these advances, our understanding of how repetitive DNA determines chromatin regulation and cellular identity is limited in aging and cancer. Although its DNA methylation landscape has been partially characterized (20), aided by the advent of long-read sequencing technologies (18), the chromatin landscape of histone post-translational modifications (HPTMs) across repetitive DNA remains largely unexplored. To address this gap, in this work we generated a large-scale atlas of six canonical HPTM signals in repetitive elements, including enhancer-associated H3K27ac and H3K4me1, promoter-associated H3K4me3, transcriptionally active H3K36me3, and repressive H3K27me3 and H3K9me3, accounting for both uniquely and multimapping reads across the BLUEPRINT ChIP-seq dataset (21). This atlas provides the most comprehensive characterization to date of cell type-specific chromatin signatures across hematopoietic lineages in repetitive DNA, and uncovers profound alterations associated with chronic hematological malignancies, as well as their interplay with chromatin changes occurring during B-cell aging.

## 2. MATERIALS AND METHODS

### 2.1. Data acquisition

Raw fastq files from ChIP-sequencing experiments targeting six canonical histone post-translational modifications (HPTMs: H3K27ac, H3K4me1, H3K4me3, H3K36me3, H3K27me3, H3K9me3), along with their respective input controls, were collected from various sorted and purified hematological cell populations and chronic lymphoid neoplasms (Fig. 1A), as provided by the BLUEPRINT epigenomics consortium (21). Our final dataset comprised 937 ChIP-seq experiments, including 766 from healthy donors and 151 from patients with hematological malignancies, obtained from the following accession codes:

**Figure 1.**
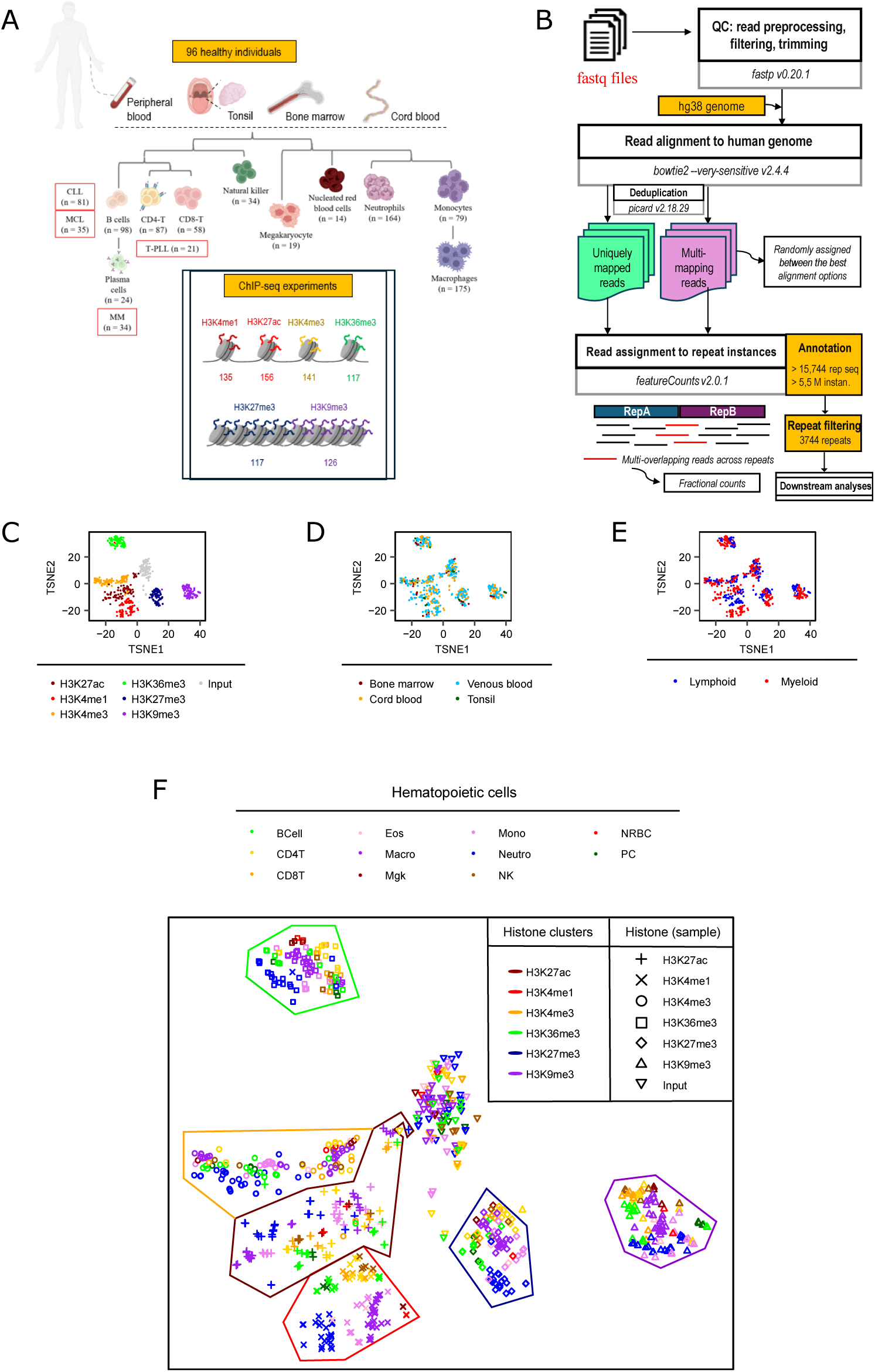
Repetitive DNA captures cell identity chromatin signatures. **(A)** Summary of the study design. **(B)** Schematic of the bioinformatic pipeline to profile chromatin signals in repetitive DNA using short-read-based ChIP sequencing experiments. **(C-F)** t-SNE dimensional reduction plots showing the distribution of samples based on the HPTM **(C)**, the hematopoietic tissue of origin **(D)**, the hematopoietic lineage **(E)**, and mature cell types **(F).**

|  |  |  |  |  |
| --- | --- | --- | --- | --- |
| EGAD00001001470, | EGAD00001001472, | EGAD00001001495, | EGAD00001001499, | EGAD00001001502, |
| EGAD00001001505, | EGAD00001001508, | EGAD00001001513, | EGAD00001001514, | EGAD00001001517, |
| EGAD00001001518, | EGAD00001001539, | EGAD00001001552, | EGAD00001001557, | EGAD00001001574, |
| EGAD00001001576, | EGAD00001001589, | EGAD00001001594, | EGAD00001002281, | EGAD00001002282, |
| EGAD00001002283, | EGAD00001002317, | EGAD00001002318, | EGAD00001002362, | EGAD00001002377, |
| EGAD00001002430, | EGAD00001002435, | EGAD00001002439, | EGAD00001002442, | EGAD00001002444, |
| EGAD00001002449, | EGAD00001002450, | EGAD00001002455, | EGAD00001002463, | EGAD00001002477, |
| EGAD00001002484, | EGAD00001002504, | EGAD00001002514, | EGAD00001002515, | EGAD00001002524, |
| EGAD00001002379, | EGAD00001002384, | EGAD00001002397, | EGAD00001002411. |  |

### 2.2. ChIP-seq data preprocessing

Single-end ChIP-sequencing reads were preprocessed using the *fastp* suite (v0.20.1) (22) with the following options: *-q 15 -l 15 --max_len1 42 --thread 9*. This enabled the removal of reads with low base Phred quality, short length (<15bp), and the trimming of potential polyG tail artifacts. Importantly, to ensure a reproducible atlas of chromatin signals in repetitive DNA based on equivalent mappability probabilities across experiments and conditions, global read trimming was applied to restrict maximum read length to 42 bp. Additionally, to account for the possibility that some experiments were encoded using Phred+64 base quality scores, *FastQC* (v0.11.9) and *fastp* (plus the *-6* option) were used in combination to detect and convert quality scores to the Phred+33 scale.

### 2.3. Genome-wide ChIP-seq profiling of HPTMs in repetitive DNA using multimapping-read-inclusive signal quantification

Processed reads were mapped to the human reference genome hg38 (initial release, obtained from: https://hgdownload.soe.ucsc.edu/goldenPath/hg38/bigZips/hg38.fa.gz) using the *bowtie2* aligner (v2.4.4) (23) with the following options: *--very-sensitive-local -p 9*. Quantification of uniquely and multimapping reads per alignment was also extracted from the *Bowtie2* output log files. The default *Bowtie2* mapping setting, which reports only the single best alignment per read or randomly selects one if multiple equally optimal alignments exist, was used, as quantification results were equivalent to those obtained when reporting potential multiple alignments (Supplementary Fig. 1A-C), while substantially reducing computational time and storage requirements (Supplementary Fig. 1D).

Next, unwanted duplicated reads were flagged and removed from coordinate-sorted BAM files with the function *picard MarkDuplicates* (v2.18.29). Subsequently, reads aligned to ENCODE blacklisted problematic ChIP-seq regions (24) were filtered out using *bedtools intersect* (v2.30.0) (25) to avoid technical artifacts, along with reads mapped to sexual chromosomes to prevent sex-based biases in downstream analyses.

Then, processed BAM files were subjected to signal quantification in repetitive DNA regions. A GTF file was generated from the corresponding RepeatMasker annotation of repetitive sequences (26), encompassing 5, 520, 118 individual instances to be analyzed in each ChIP-seq experiment. Given that random-sampling alignment strategies assign multimapping reads to one among multiple genomic loci with identical sequence, *featureCounts* (v2.0.1) (27) was used to jointly quantify uniquely and multimapping reads for each instance, and the resulting counts were subsequently collapsed across 15, 744 distinct repeat subfamilies defined by sequence within the RepeatMasker annotation. Quantification parameters included the options *--fraction* and *--largestOverlap*, which resolved reads mapping to multiple repeats by assigning them to the feature with the greatest overlap in base pairs. Finally, in cases where a read aligns to two or more repeats with identical overlap lengths, a fractional count of 1/y was assigned, where *y* represents the number of repeats involved in the multi-overlap.

### 2.4. Repeat subfamily filtering and detection of subfamilies with spurious ChIP-seq signals

Accurate detection of genomic regions exhibiting spurious signals in ChIP-seq experiments is paramount for ensuring the reliability of downstream analyses, particularly in the context of repetitive elements, some of which are enriched in known artifactual regions, such as those catalogued in the ENCODE blacklist. However, most of these regions were identified by using classical bioinformatic approaches that primarily relied on uniquely mapped reads. Given the focus of this study on repetitive DNA, we applied the following filtering criteria: (a) exclusion of repeats with low average signal (≤10 reads: 11, 568 out 15, 744 repeats, mostly microsatellites); (b) removal of repeats with an unclear sequence-based classification within the RepeatMasker annotation (125 out of 3, 906); and (c) identification of repeats with a significantly increased signal in unprecipitated chromatin, herein referred as *replist*. To define the latter, we implemented negative binomial generalized linear models (GLMs) in *DESeq2* (v1.28.1) (28), contrasting input control signals against HPTMs, while incorporating a covariate to block together matched input and ChIP experiments. This approach yielded 37 repetitive elements (FDR < 0.05 & log_2_[Input/ChIP]>0) that configure the *replist* used in the present study. Consequently, 3, 744 repeats were finally considered for downstream analyses. For visualization purposes, raw count matrices were subjected to variance-stabilizing log transformation (*vst*) according to the DESeq2 package (v1.28.1) (28).

### 2.5. Machine learning-based identification of cell type-specific chromatin signatures in repetitive DNA

Chromatin profiles of repetitive elements were initially visualized using the dimensionality reduction method *t-*distributed stochastic neighbor embedding (*t-*SNE) implemented in the *Rtsne* package (v0.16). Unsupervised clustering patterns were annotated in relation to potential sequence-related biases and diverse biological factors, including HPTMs, tissue of origin, hematopoietic lineage, and cell type.

Next, machine learning (ML) models were implemented using the *caret* package (v6.0-93) (29) to assess the predictive power of repeat-associated chromatin profiles in distinguishing terminally differentiated hematopoietic cell types, including B cells, plasma cells, CD4^+^ and CD8^+^ T cells, NK cells, neutrophils, monocytes, and macrophages. Biological replicate counts corresponding to the same HPTM, subject, and cell type were first aggregated, resulting in a matrix with 105 subject-cell type pairs and 22, 464 predictors (3, 744 repeats per 6 HTPMs) for subsequent ML analysis. Training and test sets were defined using an approximate 70:30 split across cell types, ensuring that pairs with missing data were excluded from the test set. Missing HPTM values in the training set were imputed using multiple factor analysis (MFA) implemented via the *imputeMFA* function from the *missMDA* package (v1.19) (30), which accounts for multiblock structured data during imputation.

Subsequently, learning vector quantization (LVQ) models were trained on the imputed data using 3 repetitions of 10-fold cross-validation. Cell type-specific features were identified using the *varImp* function from *caret*, retaining 2, 754 repeat-HPTM combinations with an importance score greater than 0.8 across all categories. Thereafter, multinomial logistic regression models were constructed using the function *multinom* from the *nnet* package (v7.3-14) (31), with three feature configurations: (a) the full set of 2, 754 predictors; (b) grouped by HPTM, including an input-based model as a baseline control; and (c) grouped according to RepeatMasker classes of repetitive elements. To further compare the predictive power of each repeat-annotated category at the sequence level, a total of ⌊*n/k*⌋ multinomial regression models were generated, each trained on *k* subsampled features, where ⌊⋅⌋ denotes the floor function and *k* represents the smallest number of features among repeat classes. Finally, model performance on the test set was assessed using the *predict* and *multiClassSummary* functions, and results were represented with radar plots generated by the *fmsb* package (v0.7.6).

### 2.6. Consensus-clustering discovery of chromatin epitypes in repetitive DNA

To robustly classify repetitive elements according to their chromatin landscape of HPTM, consensus clustering was performed on scaled signal intensities from 652 ChIP-seq experiments derived from healthy individuals, using the *cola* package (v2.4.0) (32). A range of clustering algorithms, including hierarchical, *k-*means, spherical *k-*means, partitioning around medoids, and model-based clustering, were applied, with the number of subgroups limited to a maximum of 10. As different numbers of consensus clusters were selected as optimal, manual inspection was employed to select a representative set of 5 chromatin epitypes with characteristic HPTM landscapes.

A limitation of module- and consensus-based clustering methods is their inability to accommodate unclassified or ambiguous features that do not fit cleanly into any cluster. To address this, repetitive elements with a Pearson correlation below 0.5 relative to the median HPTM profile of their originally inferred chromatin epitype were reassigned to a new “noise” category.

Finally, the relative importance of HPTM, cell type, hematopoietic tissue of origin, sex and age (defined as development <18 years, adult 18-55 years, and older >55 years) in explaining HPTM-specific profiles within each chromatin epitype was assessed by partitioning the total explained variance (*R*^2^) using the Lindeman, Merenda, and Gold (LMG) method, implemented in the *cal.relimp* function from the *relaimpo* package (v2.2-7) (33).

### 2.7. Chromatin state transition analyses of repetitive elements and their flaking single-copy sequences

A comprehensive full-stack chromatin state annotation for the human genome, generated with ChromHMM, was obtained from Vu and Ernst (2022) (34) to examine the relationship between the chromatin landscape of repetitive elements and surrounding single-copy sequences. Upstream and downstream flanking regions were initially defined for each repeat instance using increasing window sizes of 250bp, 500bp, 1000bp, 2000bp, and 5000bp. To retain only single-copy sequences, potential overlapping repeats within these windows were removed using the *bedtools subtract* function (v2.30.0) (25). Coordinates exhibiting artifactual signals in either repeats or flanking single-copy regions were then excluded for downstream correlative analyses. In addition, highly-similar chromatin states were consolidated into the following broad categories: *Acet* (non-enhancer-promoter acetylation), *DNase* (DNase I hypersensitivity only), *TSS* (transcription start sites), *PromF* (flanking promoters), *Tx* (transcriptional activity), *Enh* (enhancers), *BivProm* (bivalent promoters), *HET* (H3K9me3-heterochromatin), *Quies* (quiescent), *ReprPC* (H3K27me3-mediated Polycomb repression), and *znf* (zinc fingers).

To quantify the correlation between chromatin states in repeats and their corresponding flanking single-copy regions, a matrix-based state transition model (Supplementary Fig. 8A) was implemented. Within active, bivalent and repressive chromatin states, this framework assigned a maximum score of 3 to opposing states, a score of 0 to concordant states, and intermediate scores to partial or transitional state matches. Subsequently, RepeatMasker-annotated classes and families of repeats exhibiting chromatin state patterns distinct from their upstream or downstream single-copy regions were identified using Wilcoxon rank-sum tests that compared the distribution of transition scores within each category to those of elements outside the category, with effect sizes calculated using rank-biserial correlation metrics. Finally, to delineate euchromatin-heterochromatin boundaries, we reapplied our analysis framework with a modified state transition model (Supplementary Fig. 9A) that assessed the correspondence between upstream and downstream single-copy flanking regions.

### 2.8. Identification of repeat-associated HPTM alterations through negative binomial generalized linear modeling

To uncover HPTM alterations in repeats associated with chronic lymphocytic leukemia (CLL) or mantle cell lymphoma (MCL), differential analysis were conducted between each malignancy and its healthy B-cell counterpart, adjusting for sex, using the negative binomial GLM framework implemented in the *DESeq2* suite (v1.28.1) (28). This approach allowed us to account for inter-sample variability in sequencing depth and global compositional differences. Similarly, aging analyses were performed by comparing aged B-cells (>55 years) versus adult ones (18-55 yrs). *P*-values were then adjusted for multiple testing using the Benjamini-Hochberg method, and differentially enriched repeats were defined at a false discovery rate (FDR) *<* 0.05 in each comparison.

### 2.9. Identification of transitions between repetitive element epitypes

Given that multiple repeats were altered across distinct HPTMs, we sought to determine whether previously defined RE epitypes experienced transitions into different epitypes during malignant transformation. By integrating all chromatin signals in REs (whether statistically significant or not) our approach could reveal previously unrecognized, holistic changes in chromatin profiles.

To do so, multinomial regression models were first trained (using the methods described above with the package *nnet* (31)) to classify repeats within our chromatin epitypes (training set accuracy: 95.3%, test set accuracy: 95.1%). These predictors were subsequently applied to B cell, CLL and MCL samples, and chromatin transitions were defined for each disease when repeats displayed epitypes distinct from those observed in B cells.

### 2.10. Identification of regulatory networks influenced by repetitive elements

We sought to determine whether repetitive elements organized regulatory networks that influence genes associated with specific biological conditions. As genes can harbor a variable number of repeats within their annotated sequences, with longer genes exhibiting a higher probability of containing more repetitive elements, the following strategy was applied: (1) pre-selection of genes with at least 1 repeat within its sequence; (2) assignment to a specific B-CLL or B-MCL pairwise combination of chromatin epitypes when at least 50% of repetitive elements belonged to the same category, and excluded otherwise (strict filtering); (3) for each combination and condition, pathway enrichment analyses were performed using the function *enricher* from the *clusterProfiler* package (v4.6.2) (35) on those C5 gene ontologies with at least 5 and no more than 500 genes from the Molecular Signatures Database (MSigDB) (36), accessed via the *msigdbr* package (v7.5.1); (4) over-enriched gene sets were selected with FDR<0.05. For visualization, overall significant results were plotted using the UpSetR package (v1.4.0) (37).

### 2.11. Enrichment analyses of repetitive elements in enhancer regions

To determine whether significantly altered repeats were influencing the epigenetic states of distinct enhancer regions, we compiled enhancer BED annotations corresponding to major human tissues and blood cell types (38), including enhancers specific to CLL (39, 40) and MCL (41, 42), and performed permutation tests using the *regioneR* package (v1.20.1) (43). For each comparison, we quantified the number of repetitive instances overlapping enhancer regions with the *numOverlaps* function. Subsequently, to assess whether the observed overlap exceeded that expected by chance, we used the *permTest* function, generating 1, 000 randomized datasets in which the repeats’ genomic coordinates were shuffled across the human genome with the *randomizeRegions* function. The empirical significance of the observed overlap was then calculated by comparing it to the null distribution derived from these randomized datasets. Z-scores derived from the permutation tests were used to quantify the degree of enrichment of repetitive elements within enhancer regions.

### 2.12. Repeat annotation and enrichment analyses

Repetitive elements were annotated to a unique gene context using the package *annotatr* (v1.24.0) (44), When elements overlapped multiple categories, we applied the following hierarchical priority: promoter > 5’UTR > 3’UTR > first exon > exon > intron > 1-5kb upstream > intergenic. Annotations were aggregated at the subfamily level when more than 50% of the constituent repeats corresponded to the same category; otherwise, they were classified as mixed.

Regarding CpG islands (CGI), coordinates were retrieved from the UCSC *hg38* unmasked track. CpG shores (±2 kb from CGIs) and shelves (±2–4 kb) were defined accordingly, and annotations were applied with the following priority: island > shore > shelf > open sea. Subsequently, subfamilies were assigned to unique categories when 50% of repeats belonged to the same category, and labeled as mixed otherwise. Additionally, repeats for which both intergenic and intron annotations individually accounted for at least 25% of annotations were annotated as “intron/intergenic”.

Next, repeats were assigned to universal human chromatin states using the broad categories previously described. Because the number of potential annotations exceeded that of gene or CGI, chromatin states were summarized at the subfamily level by assigning the most frequent annotation when at least 25% of repeats shared it; repeats not meeting this threshold were labeled as mixed.

Finally, associations between annotations or intersections across comparisons were assessed using Fisher’s exact tests, with odds ratios (OR) serving as a measure of enrichment for a given feature.

## 3. RESULTS

### 3.1. An atlas of chromatin profiles in repetitive DNA unveils lineage and cell-type specific patterns within the human hematopoietic compartment

To uncover potential cell-type-specific determinants that shape the chromatin landscape of repetitive DNA, we compiled comprehensive genome-wide histone post-translational modification (HPTM) profiles in 766 chromatin immunoprecipitation (ChIP) sequencing experiments from the BLUEPRINT consortium (21) encompassing six well-characterized HPTMs (H3K27ac, H3K4me1, H3K4me3, H3K36me3, H3K27me3, H3K9me3) in a wide variety of healthy terminally-differentiated hematopoietic primary cells (Fig. 1A, Supplementary Table 1). While the distribution of reads across experiments was HPTM-dependent (Supplementary Fig. 2A, left), all marks contained a considerable proportion of multi-mapping reads (medians ranging from 28.08% [H3K4me3] to 46.25% [H3K9me3], Supplementary Fig. 2A, right), underscoring the magnitude of meaningful, and typically unexplored, biological information contained within the sequencing data. Indeed, while uniquely-mapping reads contributed similarly to signal at repetitive- and single-copy regions (*r*=0.67 and 0.79, respectively, both *p* < 0.001, Supplementary Fig. 2B, top), multi-mapping reads contributed substantially more to repetitive regions (*r*=0.85, *p* < 0.001) than to single-copy ones (*r*=0.43, *p* < 0.001, Supplementary Fig. 2B, bottom). Thus, aiming to interrogate chromatin signals at repetitive

DNA elements (REs), our bioinformatic pipeline (Fig. 1B, Materials and methods) integrated both uniquely mapped and classically-discarded multi-mapping reads across 5, 520, 118 individual repeat instances per sample. The main RE classes included in the annotation, when considered in terms of genome coverage (Supplementary Fig. 2C), were LINEs (41.8% of total sequence covered by REs), SINEs (25.8%), LTRs (17.5%), DNA transposons (6.7%), satellites (4.9%), and microsatellites (2.5%). Given that multi-mapping reads can be assigned by nature to multiple positions, genomic information was collapsed across individual repeats to 15, 474 independent RE subfamilies. Because RE regions may exhibit spurious ChIP-seq signal (24), we further filtered this set based on the following criteria (Materials and methods): (a) discarding REs with weak overall signal, (primarily microsatellites); (b) excluding REs with poorly defined annotation categories; and (c) removing REs with evidence of spurious signal. For this latter step, we used input libraries and negative binomial GLMs to detect REs with a significantly higher signal in unprecipitated chromatin samples, which we designated as *replist* (Supplementary Fig. 2D, Supplementary Table 2). Ultimately, the final number of selected elements was 3, 744 (24.2% of sequence-based repetitive subfamilies, most of which were classified as microsatellites [69.7%], LTRs [15.1%], DNA transposons [6.0%], and LINEs [4.6%]) and integrated the majority of the original individual genomic RE instances (5, 147, 378: 93.2% of the unfiltered annotation).

Next, we performed unsupervised exploratory analyses in our RE hematopoietic chromatin atlas to uncover the major factors driving variability using t-distributed stochastic neighbor embedding (t-SNE) techniques. First, all HPTMs displayed specific signal distributions in REs which were distinct from those of unprecipitated chromatin (Fig. 1C) and were not strongly associated with technical sequencing library factors (Supplementary Fig. 3). Within each HPTM cluster, we did not detect any noticeable grouping based on hematopoietic tissue of origin (Fig. 1D), but a clear segregation of samples based on hematopoietic lineage was observed (Fig. 1E), particularly for enhancer-like HPTMs (H3K4me1, H3K27ac). Indeed, when mapping specific cell types, we observed consistent sample grouping across the different types of hematopoietic primary cells for most HPTMs, while input controls showed no grouping capacity (Fig. 1F). These findings suggest that the epigenomic landscape of REs harbors information related to lineage differentiation.

To quantify this association, we employed artificial neural networks and multinomial regression models to predict cell of origin using the RE epigenomic data (Supplementary Fig. 4A, Materials and methods). A final set of 2, 754 HPTM-RE selected combinations (742 H3K4me1, 618 H3K27me3, 568 H3K27ac, 340 H3K4me3, 265 H3K36me3, and 221 H3K9me3) led to unsupervised segregation of cell types in both the training and test datasets (Supplementary Fig. S4B). Specifically, a multinomial predictor built with the aforementioned features showed very high levels of accuracy, sensitivity and specificity in the test set predictions (all >88%, Supplementary Fig. S4C). When stratifying features by HPTM (Supplementary Fig. S4D), we found that models based on enhancer or active HPTMs exhibited higher predictive power (mean sensitivity range: 0.78 [H3K27ac] - 0.94 [H3K4me1]]) than those based on heterochromatin marks (mean sensitivity: 0.52 [H3K27me3], 0.48 [H3K9me3]). Nonetheless, all HPTMs performed substantially better than the input control (mean sensitivity: 0.28, random expectation: 0.125). Interestingly, we observed that REs selected as predictive features in the full model were differentially distributed across HPTMs (Supplementary Fig. 4E, top). Among the most representative RE classes (all Fisher’s *p* < 0.05, all OR > 1), LTRs were enriched in H3K27ac, H3K4me1, H3K36me3, and H3K9me3 predictors; DNA transposons in H3K27ac, H3K4me1, H3K4me3, and H3K9me3; and LINEs only in H3K27me3. Microsatellites, although ubiquitously present, were not more abundant than expected relative to the reference (*p ≥* 0.05, OR < 1). Consequently, these REs spanned varying regions of the human genome (Supplementary Fig. 4E, bottom): for instance, LTRs were the dominant genomic feature in H3K27ac, H3K36me3 and H3K9me3; and LINEs predominated in H3K4me1, H3K4me3, and H3K27me3. The differential distribution of REs could reflect intrinsic, sequence-based characteristics of each RE class. To explore this, we built multinomial regression models exclusively based on individual RepeatMasker annotated classes. As expected, those RE classes with a greater number of total features exhibited improved performance, although all classes could predict above random expectation even with low features (accuracy range: 0.63 [SINE] - 0.98 [microsatellite]). Moreover, constructing models by subsampling predictors to 9 elements per model (see Materials and Methods) revealed that RE classes possess comparable average predictive powers (Supplementary Fig. 5B). Together, these results demonstrate that the epigenomic status of REs carries quantitative information capable of predicting the cell of origin across multiple HPTMs, and that while distinct RE classes are differentially represented when selecting for optimal performance, most RE classes harbor molecular information related to lineage differentiation.

Thus, we hypothesized that this tight regulation of HPTM patterns in REs could allow us to track the cell of origin in cancer, specifically, in chronic hematological malignancies (ChrHM), which partially resemble their healthy mature counterparts (45). To this end, we quantified the RE chromatin landscape of 171 additional primary chronic leukemia ChIP-seq experiments from the BLUEPRINT consortium (21) (Fig. 1A), including cases of chronic lymphocytic leukemia (CLL, B cell origin), mantle cell lymphoma (MCL, B cell origin), multiple myeloma (MM, plasma cell origin), and T-cell prolymphocytic leukemia (T-PLL, T cell origin). We then applied our previously developed multinomial regression model, trained exclusively on healthy samples, to classify these ChrHM based on their cell of origin (Supplementary Fig. 6A). When dissecting performance metrics (Supplementary Fig. 6B), the full model (followed by the H3K4me1 enhancer-mark prediction) achieved the highest mean sensitivity result on ChrHM (full: 100%, H3K4me1: 89%). Notably, all HPTMs demonstrated strong classification sensitivities (>51%), whereas input controls exhibited no predictive power (17%, random expectation: 12.5%). Interestingly, H3K4me1-based features were not uniformly modified across cell types but rather segregated into distinct clusters according to their relative levels across samples (Supplementary Fig. 6C). These clusters were partially enriched in specific RE classes (e.g., clusters 1, 4 and 5 in LTRs, cluster 2 in SINEs, cluster 3 in microsatellites, or clusters 5, 6 and 7 in DNA repeats; all Fisher’s *p* < 0.05, OR>1) but also segregated REs into epigenomic groups independently of their sequence-based classification, thus suggesting the existence of chromatin signatures specific to REs. Collectively, these findings reveal that the chromatin landscape of REs can be used to track cell lineage in chronic hematological malignancies and that distinct REs may share epigenomic profiles in a class-independent manner.

### 3.2. Systematic identification of combinatorial chromatin modification epitypes in repetitive DNA

Spatial combinations of HPTMs delineate distinct epigenomic signatures linked to genome-wide regulatory functions at single-copy DNA (46, 47). However, the specific combinatorial patterns of HPTMs present in repetitive DNA remain poorly characterized and require the inclusion of multimapping reads for comprehensive analysis. Consequently, we applied an unsupervised consensus clustering strategy (see Materials and Methods) to classify REs according to distinct chromatin epitypes characterized by specific HPTM profiles. With this approach, we were able to identify 5 major chromatin epitypes (Fig. 2A-B), four of which are broadly recognized, including constitutive heterochromatin (CH, with high levels of H3K9me3, H3K27me3 and low signals of active/enhancer marks), promoter-like (Pr, characterized by enrichment in CpG islands and elevated levels of H3K4me3), active transcription/enhancer (AE, with elevated levels of H3K36me3 and moderate H3K4me1 signals), and poised/repressed enhancer (PE, with concurrent elevated signals of H3K27me3 and H3K4me1). Interestingly, we discovered a novel chromatin epitype in repetitive DNA, termed dual heterochromatin (DH) (previously reported in poised enhancers (48)) characterized by the concurrent enrichment of H3K36me3 and H3K9me3. We further refined these clusters by assigning REs lacking clear HPTM profiles to a new noise category (Fig. 2B, Materials and Methods), primarily originating from DH and CH epitypes (Supplementary Fig. 7A). Most REs were subsequently ascribed (Fig. 2C, Supplementary Table 3) to Pr epitype (1, 407), followed by CH (631), AE (578), DH (493), Noise (371) and PE (264), though genome coverage was dominated by AE and CH epitypes (Supplementary Fig. 7B). In terms of evolutionary sequence conservation, Pr and CH contained more conserved regions (higher PhastCons scores, both Wilcoxon test *p <* 0.001), whereas Pr and DH showed signs of slower evolution rates (higher phyloP scores, Wilcoxon test *p <* 0.05 for Pr and *p <* 0.001 for DH) while CH appeared to experience neutral evolutionary patterns (Supplementary Fig. S7C).

**Figure 2.**
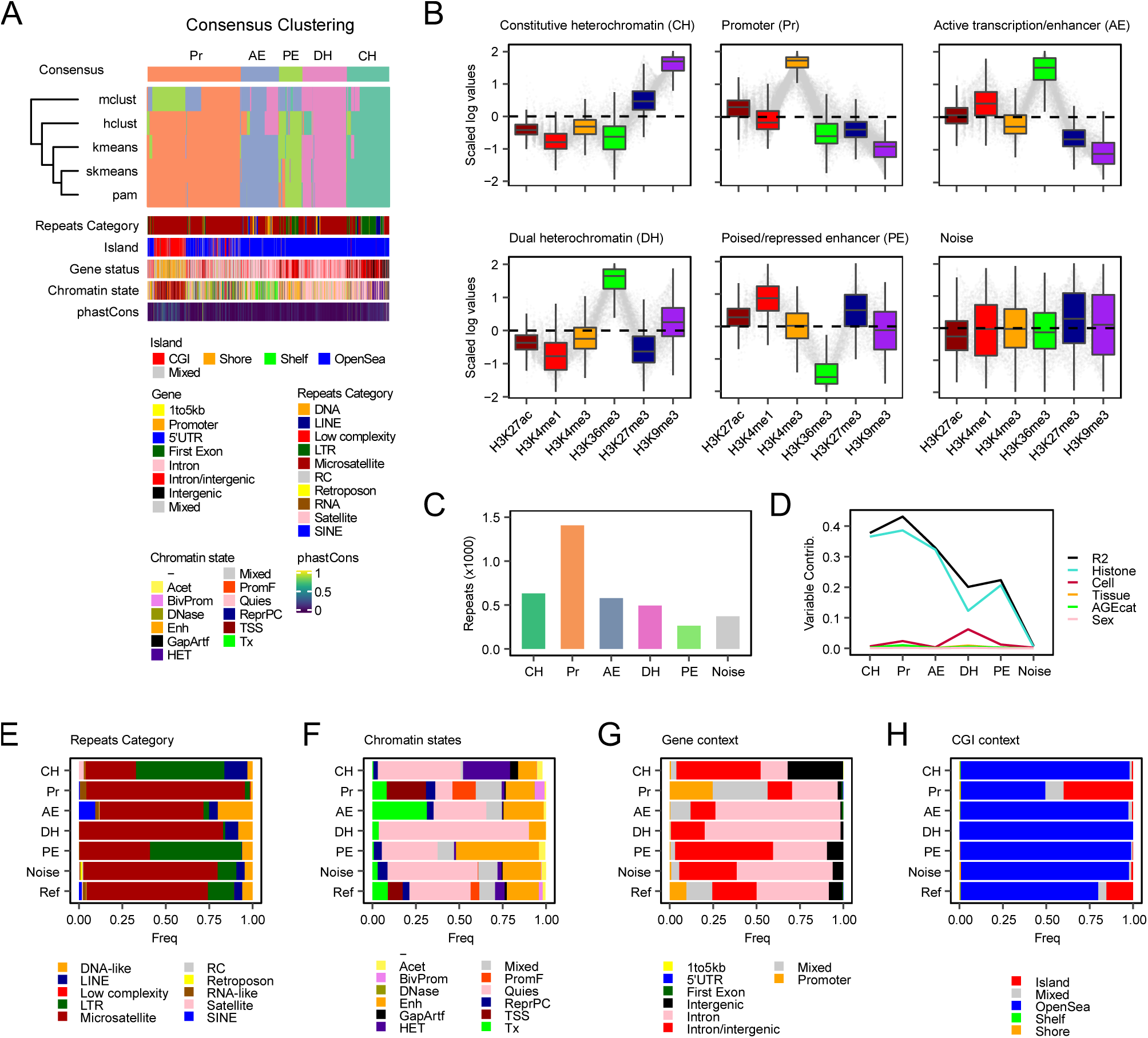
Repetitive elements adopt specific chromatin epitypes in the hematopoietic compartment. **(A)** Membership heatmap showing the results of the consensus clustering across all RE subfamilies, below, diverse genomic annotations are shown. **(B)** Boxplots showing the scaled variance-stabilizing transformed values (*vst*) of each RE subfamily, grouped by their consensus chromatin epitype. **(C)** Bar plots describing the number of RE subfamilies ascribed to each epitype. **(D)** Line plots representing absolute variable contributions in multivariate linear regression models explaining *vst* values for each chromatin epitype. Variable contributions were obtained by partitioning the total explained variance (*R^2^*) with the Lindeman, Merenda, and Gold (LMG) method. **(E-H)** Bar plots showing the relative distribution of subfamilies per chromatin epitype across sequence-based RepeatMasker-annotated classes of REs **(E)**, universal human chromatin states **(F)**, gene-based context annotations (G), and CpG island status **(H)**.

Next, we sought to explore the relative importance of biological factors underlying signal variability within each epitype. Consistent with our prior findings, histone marks were the primary contributors explaining variability across all epitypes (Fig. 2D). Sex, hematopoietic tissue of origin and age had no significant effect on the observed patterns, suggesting that chromatin epitypes of repetitive DNA are largely independent of these variables and intrinsically stable across diverse biological factors, further reinforcing their tight regulation. Importantly, cell type contribution was particularly relevant in DH (DH: 30.9%, PE: 5.5%, Pr: 5.5%), suggesting that this epitype exhibits relative plasticity within the hematopoietic compartment. As further validation, biological variables contributed negligibly to noise REs, supporting the notion of random signal.

Given the stability of chromatin epitypes in repetitive DNA across biological factors, we aimed to determine whether these patterns were associated with sequence-based features (Repeatmasker-annotated classes of REs, Fig. 2E), universal ChromHMM-derived chromatin states (based on segmentation of genomic signals from uniquely mapped reads, Fig. 2F) (34), gene context (Fig. 2G) or CpG density (Fig. 2H). No clear segregation of chromatin epitypes by specific repeat classes was observed; however, certain classes showed enrichment for certain epitypes relative to their expected frequencies in the reference (all Fisher’s *p* < 0.05, OR *>* 1), including microsatellites in Pr, LTRs in PE and CH, and SINEs and DNA transposons in AE. Similarly, major chromatin states were distributed across all epitypes, although some were clearly enriched in distinct patterns, such as flanking promoter (PromF) and transcription start sites (TSS) in Pr, heterochromatin (HET) in CH, and enhancer (Enh) in PE, consistent with biological expectations. Interestingly, our epitypes enabled us to resolve the chromatin context of most REs annotated as quiescent states, which lacked a well-defined, informative chromatin state assignment, probably due to the filtering of multi-mapping reads. On the other hand, gene context provided limited insight, except for enrichments of promoter annotations in Pr and intergenic regions in CH, while CpG islands (CGI) were enriched only in Pr, with other epitypes predominantly found in low CpG-density regions.

Together, our findings demonstrate that (1) REs can be segregated into distinct epitypes which are not primarily driven by sequence, be it at the level of RE class, genomic context or CpG density; and (2) that integrating multi-mapping reads enables the complementation of existing ChromHMM annotations, clarifying ambiguous broad annotations within repetitive regions, such as the quiescent state, which is the most prevalent.

### 3.3. LTR, Alu and SVA elements preferentially exhibit chromatin states distinct from their surrounding single-copy sequences

Given that intrinsic sequence features were not the leading determinants shaping the chromatin landscape of repetitive DNA, we sought to interrogate whether the combinatorial patterns of HPTMs in REs and their surrounding single-copy sequences exhibited significant genome-wide associations. Consequently, we analyzed chromatin states at each individual repeat locus and assessed their concordance with flanking single-copy upstream and downstream regions across varying proximity windows (Fig. 3A, see Materials and Methods). To quantify the degree of divergence, we implemented a transition model in which concordant chromatin states in REs and single-copy DNA were assigned a minimum score of 0, opposing states (e.g., active vs repressive) a maximum score of 3, and partial transitions intermediate scores (Supplementary Fig. 8A).

**Figure 3.**
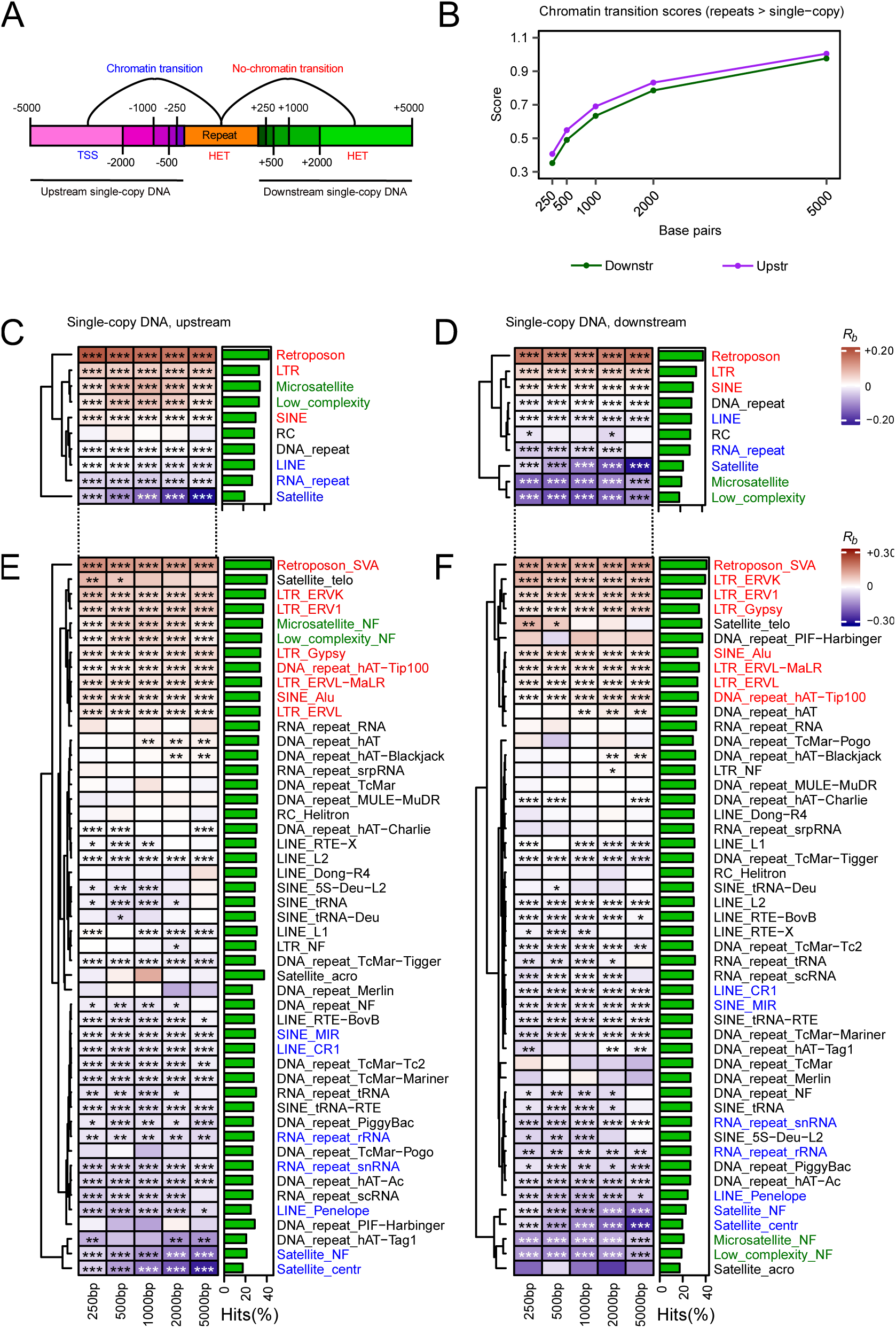
The chromatin landscape of repetitive elements and their flanking single-copy regions. **(A)** Schematic of the analysis used to compare chromatin state profiles in individual RE instances against the surrounding single-copy DNA sequences. **(B)** Line plots showing the chromatin transition scores from REs to flanking single-copy upstream (purple) and downstream (green) sequences, plotted as a function of distance (in base pairs, *bp*) from the RE. **(C-F)** Heatmaps illustrating rank-biserial correlations (*R_b_*) from Wilcoxon rank-sum tests comparing chromatin transition score vectors for each RE class/family against those of all other REs (***FDR *<* 0.001, **FDR *<* 0.01, *FDR *<* 0.05 for two-sided Wilcoxon rank sum tests). Transition scores were computed by contrasting the chromatin state of each RE instance with that of its surrounding single-copy sequences, using the values shown in Supplementary Fig. 8A. On the right side of each heatmap, bar plots indicate the average percentage of REs exhibiting any chromatin state transition across all analyzed intervals (250bp, 500bp, 1000bp, 2000bp, 5000bp). Label colors denote REs whose chromatin states are (i) distinct from both upstream and downstream surrounding single-copy regions (red), (ii) similar to both regions (blue), or (iii) similar upstream but distinct downstream (green).

First, we observed that transition scores increased genome-wide as a function of distance from REs, but, unexpectedly, were slightly more pronounced in upstream single-copy regions (Fig. 3B). Interestingly, the observed trend was largely driven by contributions from specific RE classes, such as microsatellites and retroposons, whereas satellites exhibited less asymmetric or even opposing patterns (Supplementary Fig. 8B). Subsequently, we applied Wilcoxon rank-sum tests to identify which REs transitioned more or less frequently than expected relative to their flanking single-copy DNA chromatin states (Fig. 3C-D). At the class level, LTRs, retroposons and SINEs displayed chromatin states that tend to diverge from both their surrounding upstream and downstream sequences (*p* < 0.05, *R_b_ >* 0), suggesting the involvement of specific regulatory mechanisms preferentially targeting these REs. In contrast, satellites, RNA-like REs and LINEs more frequently shared chromatin states with their flanking single-copy DNA regions (*p* < 0.05, *R_b_ >* 0). Interestingly, microsatellites and low-complexity REs showed chromatin states similar to their upstream single-copy sequences and distinct from their downstream context. Delving deeper into specific RE families (Fig. 3E-F), SVA, ERV-1/L/K and Gypsy LTRs, and Alu elements were the most distinct from their surrounding sequences, suggesting RE-specific and conserved epigenomic profiles, whereas LINE families (L1, L2, CR1, RTE and Penelope), non-Alu SINE elements and centromeric satellites were the least divergent, consistent with their heterochromatin landscape and structural roles linked to the broad H3K9me3 mark (49). In addition, we explored chromatin state transitions according to our previously defined chromatin epitypes (Supplementary Fig. 8C-D). PE, CH and AE exhibited specific chromatin states that were markedly different from those of the surrounding single-copy DNA regions, whereas DH was less prone to chromatin transitions, especially in downstream flanking regions. Interestingly, Pr elements displayed asymmetric enrichment patterns, with no continuity observed in upstream single-copy chromatin states, consistent with their colocalization at TSS regions.

Finally, in light of the observation of specific RE with asymmetric conservation of surrounding chromatin states, we investigated whether particular elements may act as potential insulators mediating the compartmentalizing of euchromatin and heterochromatin domains, based on an adaptation of our previously established chromatin transition scores (Supplementary Fig. 9A, see Materials and Methods). As expected, satellites did not exhibit such insulating function, as their extensive heterochromatin spread into adjacent regions; in contrast, LTR elements demonstrated the most pronounced genome-wide insulating behavior (Supplementary Fig. 9B). This observation was further validated at the RE family level, with Gypsy, ERV1, and ERVL LTRs identified as the most prominent elements demarcating boundaries between euchromatin and heterochromatin (Supplementary Fig. 9C).

Collectively, our results showcase how specific REs families display characteristic chromatin landscapes that differ from those of surrounding single-copy DNA sequences. Notably, SVAs, LTRs and Alu elements, along with RE epitypes marked by concurrent enrichments of H3K27me3 and H3K4me1, are the most prominent examples. Moreover, LTRs appear to exhibit the greatest potential insulator activity.

### 3.4. B-cell chronic hematological malignancies drive profound chromatin remodeling in repetitive DNA

Chronic hematological malignancies of B-cell origin, such as CLL and MCL, can harbor driver mutations in epigenetic modifiers, resulting in profound HPTM alterations at single-copy sequences (50, 51). However, studies specifically centered on repetitive DNA remain lacking. Consequently, we performed differential analyses across six HPTMs and corresponding input unprecipitated experiments between CLL/MCL samples and B cells from healthy individuals.

CLL malignancies experienced significant alterations (FDR<0.05) primarily affecting the H3K27ac mark, with 970 REs upregulated and 747 downregulated (Fig. 4A, upper left; Supplementary Table 4). In contrast, MCL neoplasms displayed widespread alterations (Fig. 4A, bottom left; Supplementary Table 5) that only excluded the Polycomb-associated H3K27me3 mark (up/down: H3K27ac 1, 297/1, 236; H3K4me1: 865/785; H3K4me3: 1, 410/1, 403; H3K36me3: 185/491; H3K9me3: 657/494). Notably, analysis of input unprecipitated controls revealed very few spurious changes in both CLL (0) and MCL (8). Although HPTM alterations in REs were directionally balanced in general, loss of HTPM signal was widespread in terms of genomic coverage across active HPTMs (Fig. 4A, right) while heterochromatin-associated H3K9me3 showed bidirectional changes in MCL.

**Figure 4.**
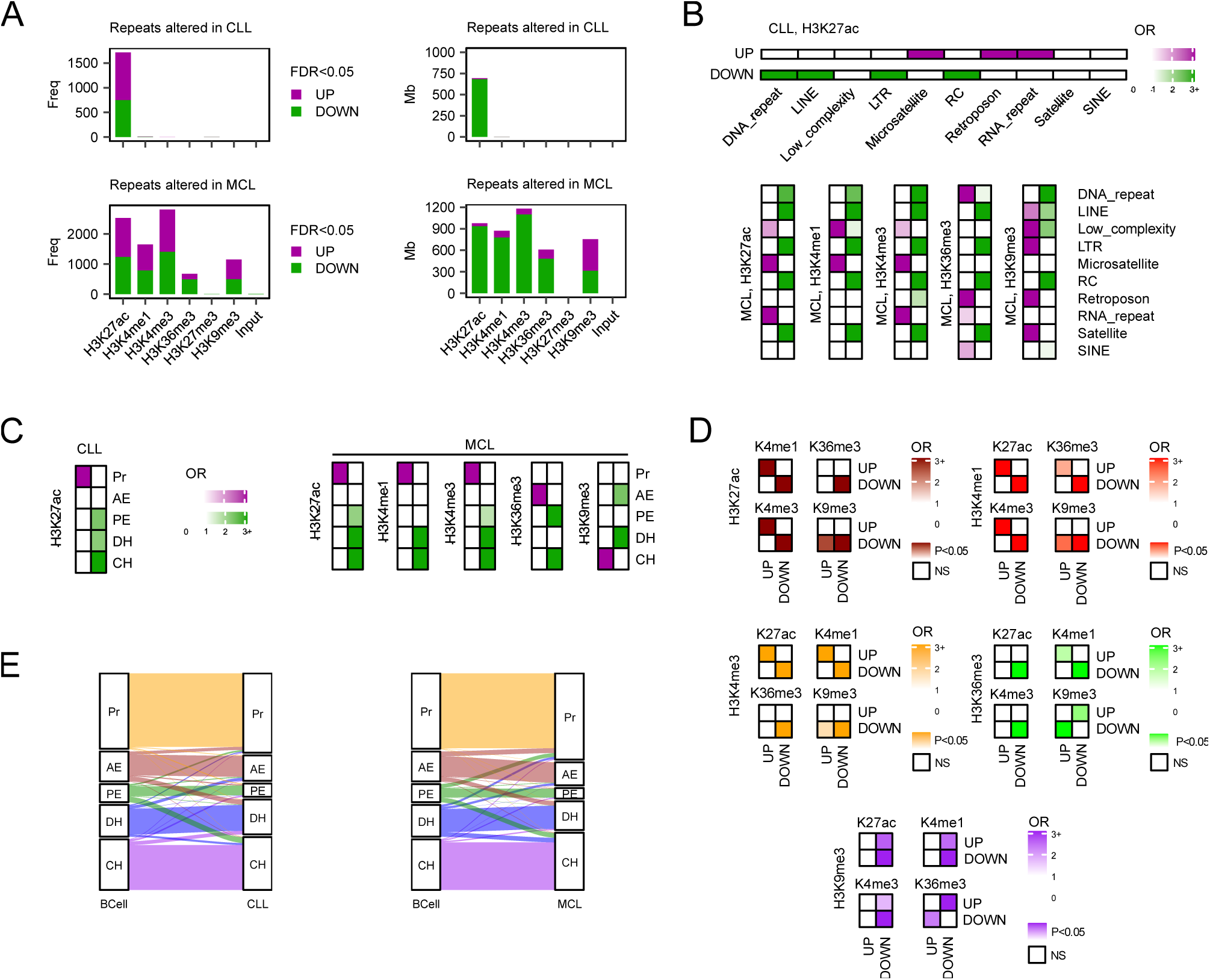
B-cell chronic malignancies reshape the chromatin landscape of repetitive DNA. **(A)** Bar plots showing the number and direction of significant (FDR<0.05) RE subfamilies (left) and their associated human genome coverage (right, Mb: mega base pairs), altered in CLL (top) and MCL (bottom) for each HPTM (upregulated: purple, downregulated: green). **(B)** Heatmaps representing the odds ratios (OR) for significantly upregulated and downregulated RE subfamilies (FDR<0.05) across RepeatMasker-annotated classes and HPTMs in CLL (top) and MCL (bottom). **(C)** Heatmaps showing the enrichment (OR) for significantly upregulated and downregulated RE subfamilies (FDR<0.05) across chromatin epitypes and HPTMs in CLL (top) and MCL (bottom). **(D)** Heatmaps depicting enrichment scores (OR) for significant intersections (Fisher’s *p<*0.05) between upregulated and downregulated RE subfamilies (FDR<0.05) across all possible HPTM pairwise comparisons. **(E)** Sankey diagrams showing the distribution of chromatin epitypes across REs and their transitions across healthy B cells and chronic malignancies (left: CLL; right: MCL) from the BLUEPRINT hematopoietic atlas.

Interestingly, these malignant HPTM alterations were not uniformly distributed across REs but were instead concentrated in specific classes (OR>1), depending on the direction of change and the type of HPTM (Fig. 4B). Microsatellites and RNA-like REs experienced gains of active chromatin marks in both B-cell malignancies, which is particularly noteworthy given that these classes are frequently engaged in active transcriptional mechanisms (e.g., TSS, rRNAs) (52, 53). This also accounted for the relatively narrow genomic span of regions acquiring active marks, as these REs occupy smaller regions of the genome (Supplementary Fig. 2C). In contrast, highly-abundant DNA-like, LINE and LTR REs, which cover large portions of the genome, exhibited a widespread loss of active marks in both neoplasms (Supplementary Fig. 10A-B), which could passively result from malignant replicative processes that impair epigenetic fidelity mechanisms and cause loss of epigenetic information (54).

Importantly, we found targeted gains of repressive H3K9me3 in MCL affecting key RE classes such as LTRs and satellites, highlighting the malignant cell’s need to maintain these sequences in an inactive state (Fig. 4B). Regarding LINEs, we observed bidirectional enrichments of H3K9me3 in MCL, with gains in L1 contrasted by losses across other families (Supplementary Fig. 10B), reflecting complex regulation that specifically repressed L1, while other non-autonomous LINE sequences underwent loss of all HPTMs. Similarly, DNA-like REs showed widespread loss of HPTMs in MCL, including H3K9me3, whereas certain families (hAT and TcMar) accounted for the targeted gains observed in H3K36me3. Remarkably, SVA retroposons acquired both H3K36me3 and H3K9me3, alterations that, when concurrent, could facilitate the formation of poised chromatin states (Supplementary Fig. 10B).

Inspired by these results, we sought to determine whether these coordinated HPTM alterations targeted specific repetitive chromatin epitypes (Fig. 4C). Consistent with our previous RE class enrichments, gains of active histone marks were concentrated on Pr epitypes, while losses in MCL were found on heterochromatin domains (CH, DH) and, to a lesser extent, in PE. Interestingly, bidirectional H3K9me3 alterations affected different heterochromatin epitypes, with gains especially targeting CH and losses DH. Regarding enhancer epitypes, significantly altered PE REs appeared to reinforce their repressing state by selectively losing active HPTMs, while AE alterations globally acquired H3K36me3 and/or reduced H3K9me3 signals in MCL. Together, these findings showcase how (1) the epigenomic landscape of REs is broadly altered in hematological malignancies, which display widespread loss of active chromatin and bidirectional changes in heterochromatin; and (2) epigenomic alterations in RE specifically target distinct classes and families which can be linked to diverse functional roles and are consistent with previously defined RE epitypes.

Depending on specific HPTM combinations, malignant alterations affecting REs can occur synergistically or antagonistically, as has been shown for single-copy sequences (55). To test this hypothesis, we systematically analyzed the overlap between REs significantly altered across distinct histone marks in MCL (Fig. 4D). Active HPTMs and H3K4me1 enhancer signals experienced synergistic alterations, while their reductions coincided with a global, nonspecific loss of both active and repressive HPTMs. Interestingly, loss of active HTPMs was also linked to gains in H3K9me3, suggesting the heterochromatinization of REs. Notably, H3K36me3 and H3K9me3 alterations were clearly antagonistic, indicating a chromatin switch between these two HPTMs.

In light of these observations, we set out to assess whether the different interactions between alterations in RE HPTMs could induce a switch in RE chromatin epitypes or merely amplify or attenuate pre-existing chromatin signals. To do so, we developed machine learning classifiers to assign REs to our 5 main chromatin epitypes based on their HPTM signals (see Materials and Methods), which allowed us to explore specific epìtype transitions occurring in healthy B cells and their potential transformations into malignant counterparts (Fig. 4E). Notably, we observed a limited number of shifts (avg: 20.1%; Pr: 2.4%; AE: 28.2%; PE: 44.6%; CH: 8.4%; DH: 16.9%) consistent with the finding that significant alterations were limited to a single HPTM, H3K27ac. Specifically, certain REs from enhancer epitypes such as PE (30.3%) and AE (15.3%) appeared to be repressed, respectively, into CH and DH domains, in agreement with loss of active HPTMs. Interestingly, a subset of poised DH and PE REs also transitioned toward activation, switching into AE (9.3%) and Pr epitypes (9.2%), respectively, albeit at lower frequencies. Regarding MCL, despite observing widespread alterations across distinct HPTMs, the overall chromatin epitype landscape remained relatively stable (avg: 22.3%; Pr: 0.6%; AE: 30.6%; PE: 60%; CH: 4.3%; DH: 25.0%), reinforcing the idea that HPTM alterations in REs across B-cell ChrHM primarily amplify or attenuate pre-existing chromatin signals. However, it is worth noting that: (1) ∼30% of B-cell-annotated AE REs transitioned into different epitypes, including repression toward DH (14.1%) and further activation toward Pr (15.1%); (2) ∼49% of B-cell-annotated PE REs shifted into CH (25.1%) or Pr (22.7%); and (3) 12.5% of B-cell-annotated DH REs were further repressed toward CH, whereas another 12.5% shifted toward activation in Pr (6.3%) and AE (6.3%) epitypes.

Altogether, the integration of all chromatin signals revealed a complex scenario of interactions between HPTMs, providing insight on how loss of active chromatin modifications can lead to heterochromatinization of REs, and the existence of an antagonistic relationship between H3K36me3 and H3K9me3, which may underlie how B-cell H3K36me3-enriched epitypes (AE, DH) transitioned into one another upon malignant transformation.

### 3.5. Repetitive DNA modulates critical oncogenic pathways through chromatin remodeling in B-cell malignancies

Having identified important alterations in the chromatin landscape of REs during B-cell malignant transformation, we next investigated whether these REs could be linked to pathways relevant to tumorigenesis. Initially, we performed overrepresentation analyses (ORA) to determine the most functionally relevant epitype transitions (see Materials and Methods). Activating switches (AE to Pr in CLL, and DH to Pr in MCL) represented the largest proportion of significant ontologies (FDR<0.05) across all transitions (Supplementary Fig. 11A). Delving deeper into their associated functions, we observed the following: (1) in CLL, AE REs enriched in pathways related to lymphocyte chemotaxis and stem cell division were further activated in Pr chromatin epitypes (Supplementary Fig. 11B); (2) in MCL, DH REs transitions to Pr preferentially targeted pathways of B-cell proliferation and immune activation (Supplementary Fig. 11C); and (3) Pr, CH, and PE switches, along with AE/DH transitions to repressive states, were not consistently enriched in particular gene ontologies.

Notably, Pr REs that did not undergo any transition showed the highest number of significantly enriched pathways (Supplementary Fig. 11A), mainly related to embryonic and developmental processes (Supplementary Fig. 11D). This finding aligns with the preservation of the differentiated B-cell-like state observed in both CLL and MCL, in contrast to other acute leukemia that display a more stem-like or undifferentiated phenotype. Similarly, AE REs associated with core biosynthetic pathways such as gene expression, splicing, and translation, remained unaltered in healthy B cells and chronic malignancies (Supplementary Fig. 11D). Additionally, unmodified CH REs were enriched in pathways unrelated to both the pathophysiology of these diseases and the biology of B cells (Supplementary Fig. 11D), consistent with constitutive mechanisms of H3K9me3-mediated heterochromatin repression. Interestingly, poised DH REs shared by CLL and healthy B cells were found to be enriched in B-cell proliferation and immune activation pathways that were upregulated in MCL, in line with its more proliferative and aggressive phenotype relative to CLL (Supplementary Fig. 11D). All in all, chromatin activating transitions from B cells revealed the upregulation of pathways relevant to the pathophysiology of CLL and MCL malignancies, while embryonic gene ontologies remained mostly unaffected.

Finally, to further understand the regulatory network under RE control, we investigated the impact of H3K27ac alterations on enhancers relevant to the pathophysiology of chronic B-cell malignancies. In CLL (Supplementary Fig. 12A), REs exhibiting gains of this active enhancer HPTM (FDR<0.05, log2(FC)>1) were markedly enriched at CD34+ enhancers, which are characteristic of hematopoietic stem and progenitor cells (HSPCs). Similarly, in MCL, H3K27ac-gaining REs were predominantly concentrated at CD34+ enhancers (Supplementary Fig. 12B), whereas REs showing loss of this HPTM were largely underrepresented at these enhancers (Supplementary Fig. 12C). Additionally, investigating other HPTMs which were also substantially altered in MCL revealed significant enrichments at CD34+ enhancers for H3K4me3, and, to a lesser extent, H3K4me1 (Supplementary Fig. 12D). Altogether, our findings provide compelling evidence that REs might contribute to the epigenetic activation of stem cell-like enhancers in chronic malignancies of B-cell origin.

### 3.6. Aging in the B-cell compartment primes the epigenetic landscape of REs to carcinogenic transformation

Age-associated alterations across distinct molecular layers have been consistently linked to the erosion of protective mechanisms that ultimately predispose to oncogenesis (56, 57). However, the specific epigenetic relevance of repetitive DNA remains unresolved. Consequently, we performed comprehensive epigenomic profiling of HPTM alterations in REs across the adult-to-elderly transition in primary B cells.

Initially, we observed that chromatin remodeling during B-cell aging selectively targets heterochromatin marks (Fig. 5A, left; H3K27me3: 313 upregulated, 226 downregulated; H3K9me3: 103 upregulated, 61 downregulated; Supplementary Table 6). Although most REs were upregulated, when considering genomic coverage, 56.7% of the repetitive regions affected in H3K27me3 lost the Polycomb-associated repressive mark, whereas the vast majority of H3K9me3-altered regions (80.9%) lost constitutive heterochromatin (Fig. 5A, right), which is consistent with previously reported loss of heterochromatin during aging (58). At the level of RE class, we found that LTRs and satellites experienced targeted heterochromatinization during B-cell aging (OR>1, Fig. 5B, top), mirroring alterations observed in B-cell chronic malignancies. Conversely, LINE REs underwent passive loss of both H3K27me3 and H3K9me3, whereas DNA-like REs were selectively depleted of H3K27me3. When considering epitypes, loss of heterochromatin affected poised DH states, whereas CH and PE showed concurrent gains and losses of both repressive HPTMs, resulting in bivalent enrichments (Fig. 5B, bottom).

**Figure 5.**
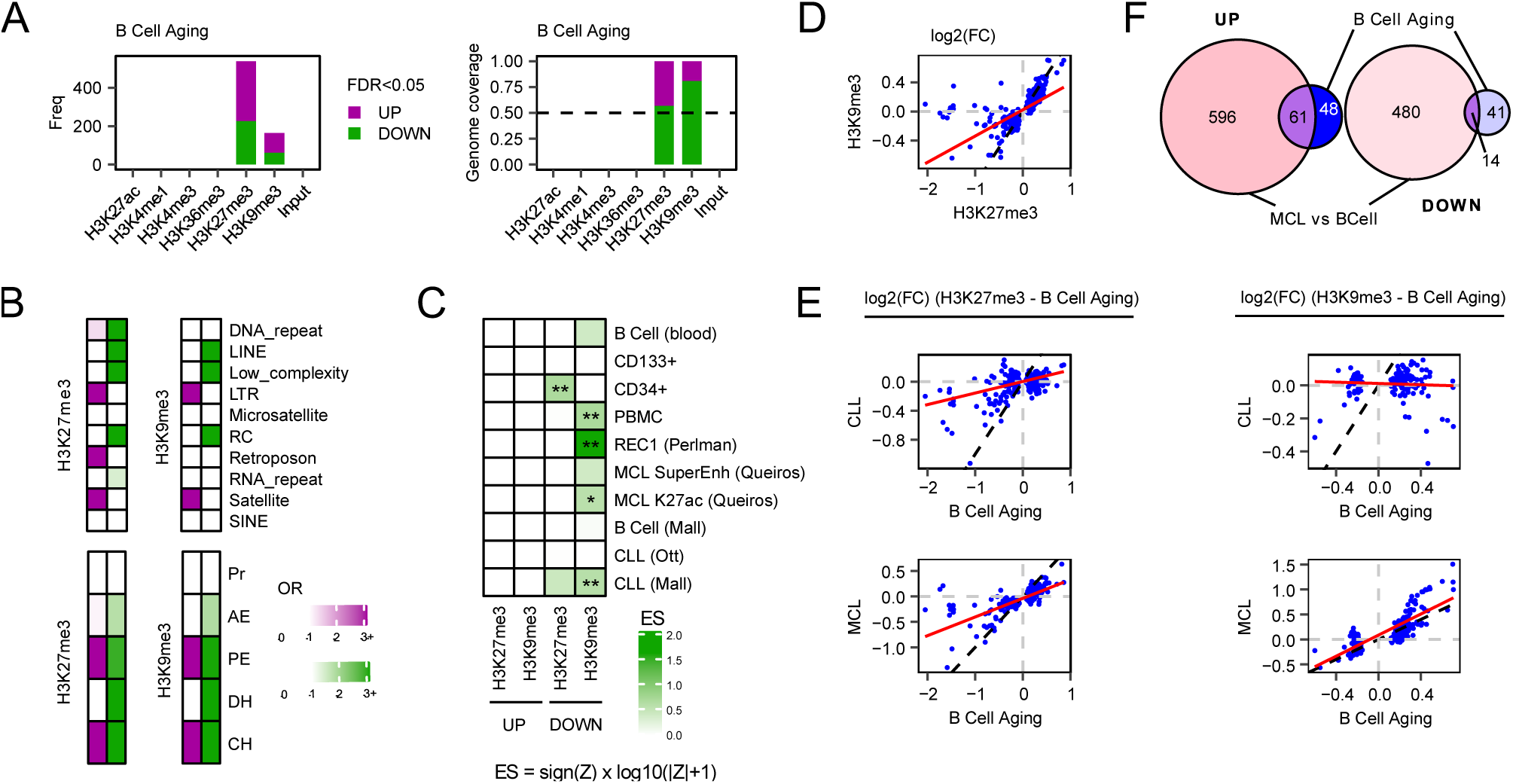
B-cell aging predisposes the chromatin landscape of repetitive DNA to malignant transformation. **(A)** Bar plots showing the number and direction of significant (FDR<0.05) RE subfamilies (left) and their associated human genome coverage (right, Mb: mega base pairs), altered during B-cell aging for each HPTM (upregulated: purple, downregulated: green). **(B)** Heatmaps representing the enrichment scores (OR) for significantly upregulated and downregulated RE subfamilies (FDR<0.05) across RepeatMasker-annotated classes (top) and chromatin epitypes (bottom) per HPTM. **(C)** Heatmap of enrichment scores (sign(Z) × log_10_(|Z|+1), where Z is the z-score obtained for one-sided permutation *regioneR* tests) depicting overlaps between distinct enhancer regions and the 50 RE subfamilies with the largest log_2_ fold changes (FDR<0.05) per HPTM, stratified by direction of change (***FDR *<* 0.001, **FDR *<* 0.01, *FDR *<* 0.05 for *regioneR* tests) **(D)** Scatter plot showing the correlation between the log_2_ fold changes for H3K27me3 versus H3K9me3 levels in REs altered in at least one of these HPTMs during B-cell aging. **(E)** Scatter plots illustrating the correlation between the log_2_ fold changes for H3K27me3 (left) and for H3K9me3 (right) in B-cell aging versus CLL (top) or MCL (bottom) across REs altered in aging for each HPTM. **(F)** Venn diagram showing the intersection between REs exhibiting significant H3K9me3 alterations (FDR<0.05) in B-cell aging and MCL, separated by direction of change.

Notably, repetitive regions losing H3K9me3 were significantly enriched at enhancers activated in CLL and MCL, and those depleted of H3K27me3 specifically targeted stem cell-like CD34+ enhancers (Fig. 5C). These results support the notion that loss of epigenetic repressive mechanisms in REs during B-cell aging may prime cells to oncogenic progression. In this line, we also found a strong trend in which gains and losses of H3K27me3 and/or H3K9me3 were correlated and occurred concurrently in REs (Fig. 5D: *r =* 0.6368, *p <* 0.001), suggesting that these HPTMs act in a concerted manner during age-associated alterations, which strengthens the priming toward malignancy in heterochromatin.

Motivated by our previous observations, we sought to explore the extent of the interplay between cancer- and age-associated alterations in REs across distinct HPTMs. First, we observed that RE heterochromatin marks significantly altered during B-cell aging showcased similar patterns in CLL (H3K27me3: *r =* 0.466, *p <* 0.001; H3K9me3: *r = –*0.065, *p =* 0.411) and MCL (H3K27me3: *r =* 0.743, *p <* 0.001; H3K9me3: *r =* 0.798, *p <* 0.001) (Fig. 5E). Additionally, MCL and B-cell aging alterations in H3K9me3 within REs (FDR<0.05) were consistently preserved (Fig. 5F; UP: OR *=* 6.48, *p <* 0.001; DOWN: OR *=* 2.28, *p =* 0.014).

Next, to gain a global view of the aging-cancer interactions in HPTM levels, we examined genome-wide correlation patterns between cancer and aging across all REs and HPTMs, irrespective of statistical significance (Supplementary Fig. 13A). In both CLL and MCL, most HPTMs displayed modest positive or neutral correlations with aging for the majority of HPTMs, whereas the active modifications H3K27ac and H3K4me3 showed strikingly strong anticorrelations. These findings indicate that malignant transformation triggers *de novo* activation of REs normally repressed during aging, a process that appears to occur independently of the aforementioned concordant changes observed in heterochromatin HPTMs. To delve deeper into this phenomenon, we explored enhancer enrichments across all HPTMs in aging and cancer. During B-cell aging, repetitive regions significantly enriched at enhancers underwent a widespread, non-specific loss of HPTMs (Supplementary Fig. 13B). Conversely, although CLL and MCL displayed an independent depletion of H3K9me3, both diseases exhibited an increased deposition of active HPTM in REs associated with enhancer regions (Supplementary Fig. 13B). All in all, our findings revealed that the heterochromatin erosion that occurs during B-cell aging, which globally affects REs, is necessary but not sufficient for the development of malignancies, which also require the targeted acquisition of active HPTMs at these primed enhancers.

## 4. DISCUSSION

Combinatorial patterns of histone post-translational modifications constitute chromatin codes that critically shape dynamic processes like hematopoietic lineage commitment (59–61). Yet most current insights come from ChIP-seq analyses restricted to single-copy genomic regions, which exclude multimapping reads and limit characterization of chromatin states within repetitive DNA (16). Here, we generated a comprehensive atlas of repetitive-DNA chromatin profiles across the human hematopoietic compartment by integrating multimapping reads from 937 ChIP-seq datasets for six well-known HPTMs (H3K27ac, H3K4me1, H3K4me3, H3K36me3, H3K27me3, H3K9me3) into 3, 744 repeat subfamilies with robust biological signals.

In healthy, mature hematopoietic cells, we found that repetitive-DNA chromatin signals are HPTM-specific and reliably track hematopoietic lineage, highlighting the epigenetic relevance of the dark genome (62). Despite substantial sequence divergence, all RE classes retained cell identity information, paralleling the specificity of DNA methylation reported for mouse REs (63). Inspired by these findings, we hypothesized that repetitive DNA harbors specific chromatin codes, leading to the discovery of 5 major RE epitypes: promoter-like active regions, constitutive heterochromatin, poised or active enhancers, and a dual heterochromatin state marked by concurrent H3K36me3 and H3K9me3, previously reported at poised cell-identity enhancers (64), which exhibited the greatest cellular plasticity among epitypes. Our annotations also aligned with chromatin states derived from uniquely mapping reads (34) and enabled finer classification of RE epigenomic states, clarifying elements previously labeled quiescent. Collectively, our findings demonstrate that REs preserve defined epigenomic features, reflecting complex, context-dependent regulation, and revealing patterns consistent with the epigenetic control of specific RE families (65), chromosomal regions (66), and global chromatin signals (67).

Interestingly, chromatin epitypes were not primarily driven by sequence features, prompting us to examine the interplay between chromatin states in REs and surrounding single-copy regions. By analyzing chromatin state transitions, we identified ERV/Gypsy LTRs, SVA and Alu sequences as the most epigenetically distinct, suggesting specialized regulation relative to single-copy DNA. These findings are particularly relevant given that LTR can regulate genes *in trans* (68–70), Alu elements orchestrate long-range enhancer-promoter interactions (71), and SVA transcripts drive long-distance activation of genes controlling myelopoiesis and erythropoiesis (72). Notably, we observed that LTRs display insulator activity broader than that previously reported for subsets of MIRs (73) and tRNAs (74), though it remains to be determined whether they can block enhancer function beyond acting as chromatin domain barriers. In contrast, we found that satellite and LINE REs are largely epigenetically similar to their surrounding single-copy regions, consistent with centromere position being defined primarily by heterochromatin boundaries rather than satellite sequence (75), and in line with observations that L1 elements contribute to transcriptomic neuronal networks via bidirectional promoters (76).

After observing that REs form functional compartments, with particular families preserving specific chromatin features, we investigated how repetitive epigenomes are targeted in two interrelated processes, cancer and aging. Within the B-cell compartment, we detected extensive HPTM alterations in MCL and CLL-specific disruptions of H3K27ac, consistent with previous studies focused on single-copy sequences (50). Gains and losses of HPTMs were coordinated across chromatin epitypes and RE classes, reflecting structured, element-specific regulatory programs linked to malignant transformations. Active HPTMs were preferentially lost across RE families that gain H3K9me3, including LTR, L1 and satellite elements, suggesting targeted heterochromatinization, whereas other families displayed non-specific loss of both repressive and active HPTMs, potentially reflecting impaired epigenetic fidelity during mitosis.

Collectively, our findings indicate that heterochromatin remodeling in tumorigenesis is governed by regulatory dynamics far more complex than previously recognized in repetitive DNA. Prior studies linked loss of H3K9me3 to increased genomic instability, including aberrant chromosomal translocations, structural abnormalities, and illegitimate recombination of repetitive elements (77–79), whereas heterochromatin spreading associates with cell differentiation (80), phenotypic transcriptional plasticity (81), and elevated mutation rates (82, 83). Interestingly, heterochromatin disruption in lymphoma has been found to perturb B cell transcriptional networks and trigger cytotoxic responses (84), a phenomenon also observed in stem cells from chronic malignancies through transposon activation (85). Furthermore, heterochromatin erosion and subsequent transposon expression have been reported to protect against solid tumor development through innate immune activation (86–88), while contributing to increased inflammatory states during aging (89). We therefore hypothesize that transposon transcriptional activation via epigenetic rewiring may represent a novel therapeutic vulnerability in chronic malignancies. Altogether, our findings support a model in which heterochromatinization occurs across satellite repeats, potentially facilitating telomere elongation and cell replication (90), and across LTR/L1 transposon elements, where it may restrain innate immunogenicity (86–88).

On the other hand, although repetitive DNA broadly lost active HPTMs, targeted gains occurred in promoter-like REs and microsatellites. Strikingly, genes with REs that transitioned to promoter-like epitypes were highly enriched in pathways central to malignant pathophysiology, including B-cell proliferation, stem-cell division, immune activation, and chemotaxis. Furthermore, H3K27ac alterations in REs were linked to aberrant activation of CD34+ stem-cell enhancers. Finally, we also uncovered a chromatin switch connecting H3K36me3 and H3K9me3, in which increases in one modification were tightly coupled to decreases in the other. This phenomenon was associated with malignant reprogramming of chromatin epitypes and may drive interconversion between dual heterochromatin and active-enhancer repetitive regions. Of note is the fact that SVA elements showed co-enrichment of both H3K36me3 and H3K9me3, consistent with recent evidence demonstrating that activation of bivalent SVAs induces myelopoietic programs normally silenced in the lymphoid lineage (72).

Altogether, our findings provide compelling evidence that repetitive DNA plays a major, previously unrecognized role in remodeling gene regulatory networks that drive malignant progression within the hematopoietic compartment. Hence, we hypothesized that aged B cells may acquire chromatin changes at their REs that prime them for malignant development. Consistent with the loss-of-heterochromatin model of aging (91), we initially observed a global erosion of H3K9me3 patterns in repetitive DNA. Moreover, these alterations were not offset by changes in H3K27me3, a compensatory mechanism previously described in single-copy sequences (55), and are consistent with a model of coordinated behavior involving H3K27me3 and H3K9me3 in repetitive DNA, as suggested by prior studies (92, 93). Interestingly, we also detected gains of heterochromatin marks targeting LTRs and satellites, mirroring the chromatin alterations observed in B-cell malignancies. Therefore, it seems that REs may accumulate epigenomic damage during aging that can be partially traced to malignant transformation. Notably, heterochromatin loss at REs during B-cell aging also targets regulatory sequences, which may represent an early priming event preceding malignant transformation. In fact, our analyses revealed that CLL and MCL specific enhancers are enriched at H3K9me3-depleted REs, whereas CD34+ stem cell enhancers are associated with REs exhibiting H3K27me3 loss. Conversely, we also observed strong global anticorrelations between age-associated and tumorigenic dynamics in active HPTMs, suggesting *de novo* activation in cancer (50) mediated by REs at these regulatory sequences.

In summary, we have provided a foundational resource for understanding how chromatin dynamics at repetitive DNA influence hematopoietic lineage commitment, aging and malignant transformation within the human B-cell compartment.

## Acknowledgements

We sincerely thank the BLUEPRINT Epigenomic Consortium for generating high-quality raw sequencing data that were instrumental to this study. We also acknowledge access to and expert technical support from the SCAYLE Supercomputing Center (León, Spain), whose high-performance computing infrastructure enabled efficient data processing and large-scale storage.

## Author contributions

JJAL, RFP and MFF conceived, coordinated and supervised the study. JJAL designed and performed the main data analyses. JRT, LSL and AFF assisted with the interpretation of results. JJAL, RFP and MFF participated in drafting the manuscript. All authors revised and approved the final manuscript.

## Data availability

The sequencing data underlying this article are available under restricted access in accordance with BLUEPRINT authorization rules, and corresponding accession codes are provided in the Materials and Methods section.

## Supplementary figures

**Supplementary Fig. 1.**
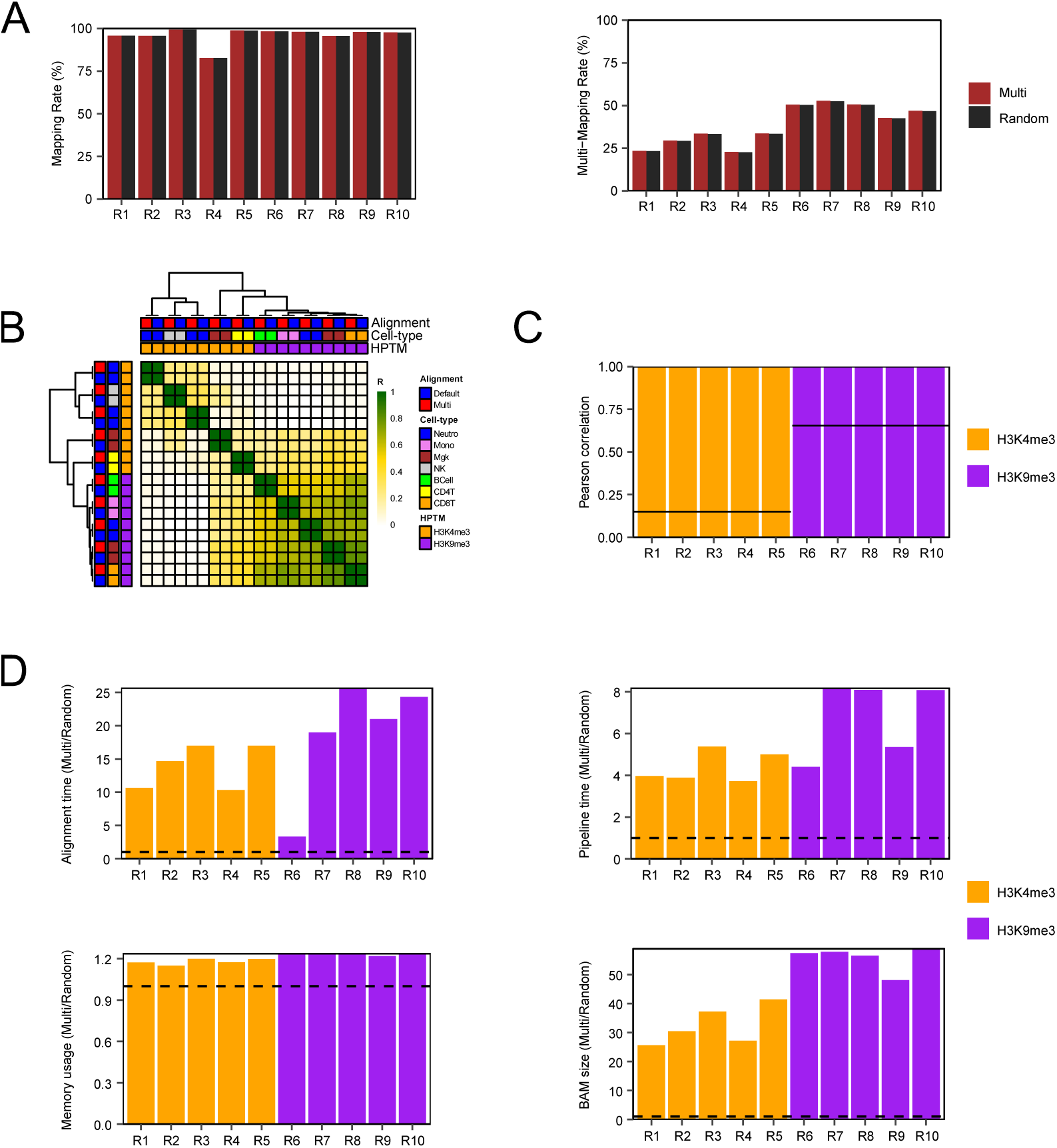
Comparison of alternative alignment pipelines for quantifying ChIP-seq multimapping reads in repetitive DNA. (A) Bar plots showing the concordance of ChIP-seq mapping and multimapping rates across 10 sequencing experiments of 100k reads, analyzed using two different alignment modes to the hg38 human genome: (1) random allocation of multi-mapping reads (Random, in black); (2) allocation of multi-mapping reads for up to 1, 000 valid alignments (Multi, *bowtie2 -k 1000*, in red) and assignment of fractional counts. (B) Heatmap illustrating the high correlation between ChIP-seq counts ascribed to REs through the Random or Multi pipeline. (C) Bar plots depicting the pairwise Multi/Random Pearson correlation of RE counts per sequencing experiment and HPTM (H3K4me3: yellow, H3K9me3: purple), with the expected average inter-sample correlation per HPTM indicated by horizontal lines. (D) Bar plots showing the Multi/Random ratio per HPTM (H3K4me3: yellow, H3K9me3: purple) for alignment and pipeline runtime, memory usage, and BAM file size, demonstrating that Random pipeline produces equivalent results across all sequencing experiments while substantially reducing computational time and resource usage.

**Supplementary Fig. 2.**
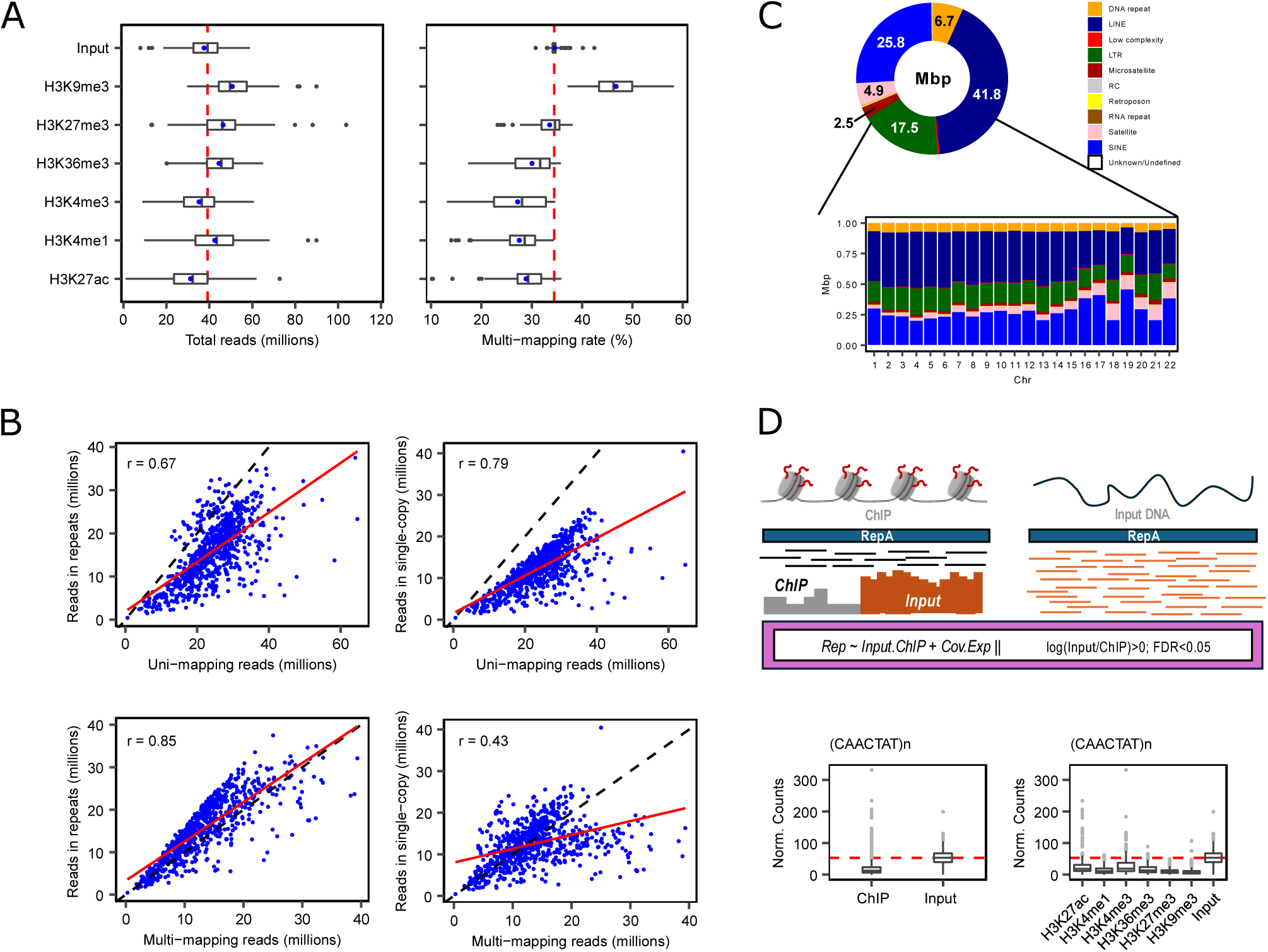
Overview of repetitive DNA preliminary analyses applied to ChIP-seq hematopoietic data. (A) On the left, boxplots depicting the distribution of ChIP sequencing reads per histone mark in healthy hematopoietic cell types from the BLUEPRINT consortium. On the right, boxplots showing the percentage of multi-mapping reads over the total number of reads. Red dashed lines represent in both cases the median values for input experiments. (B) Scatter plots showing the linear correlation (in red) between reads annotated to REs and uni-mapping (top left) or multi-mapping reads (bottom left) and between reads annotated to single-copy regions and uni-mapping (top right) or multi-mapping reads (bottom right). Perfect linear correlation is represented using black dashed lines. (C) Above, donut plot showing the percentual distribution of main repeat categories across human hg38 annotation (Mbp: mega base pairs). Below, bar plots depicting the percentual distribution of main repeat categories per autosomal chromosomes. (D) Above, schematic of the computational approach to detect repeats with a spurious background signal (*replist* elements). Below, boxplots showing the normalized counts of a *replist* element across ChIP/Input experiments (left) and segregated by HPTM (right).

**Supplementary Fig. 3.**
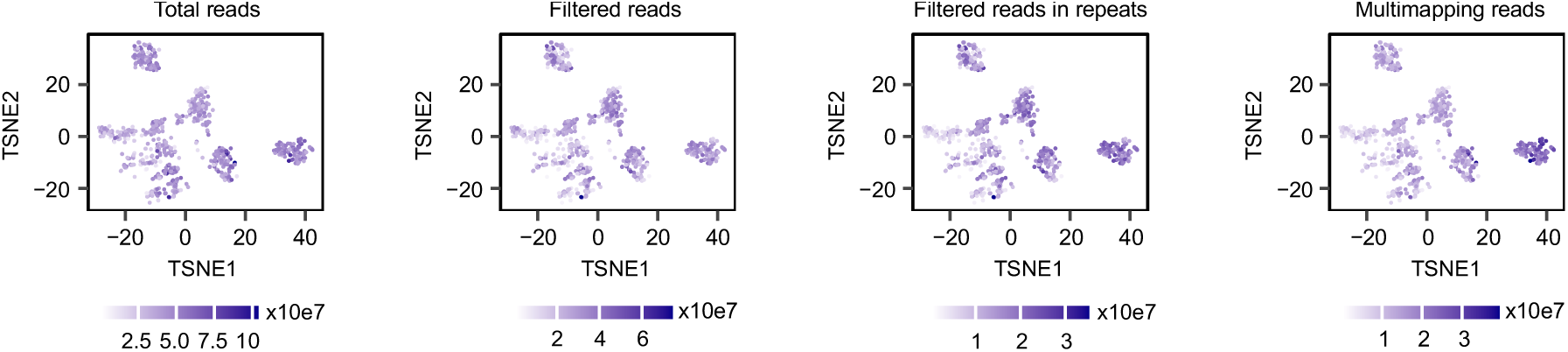
Exploring the repetitive DNA chromatin landscape across hematopoietic samples using sequencing-based metrics. t-SNE-based dimensional reduction plots showing the distribution of samples according to the total number of ChIP-seq reads (left), reads retained after different filtering steps (middle left), those mapping to REs (middle right), and multi-mapping reads (right).

**Supplementary Fig. 4.**
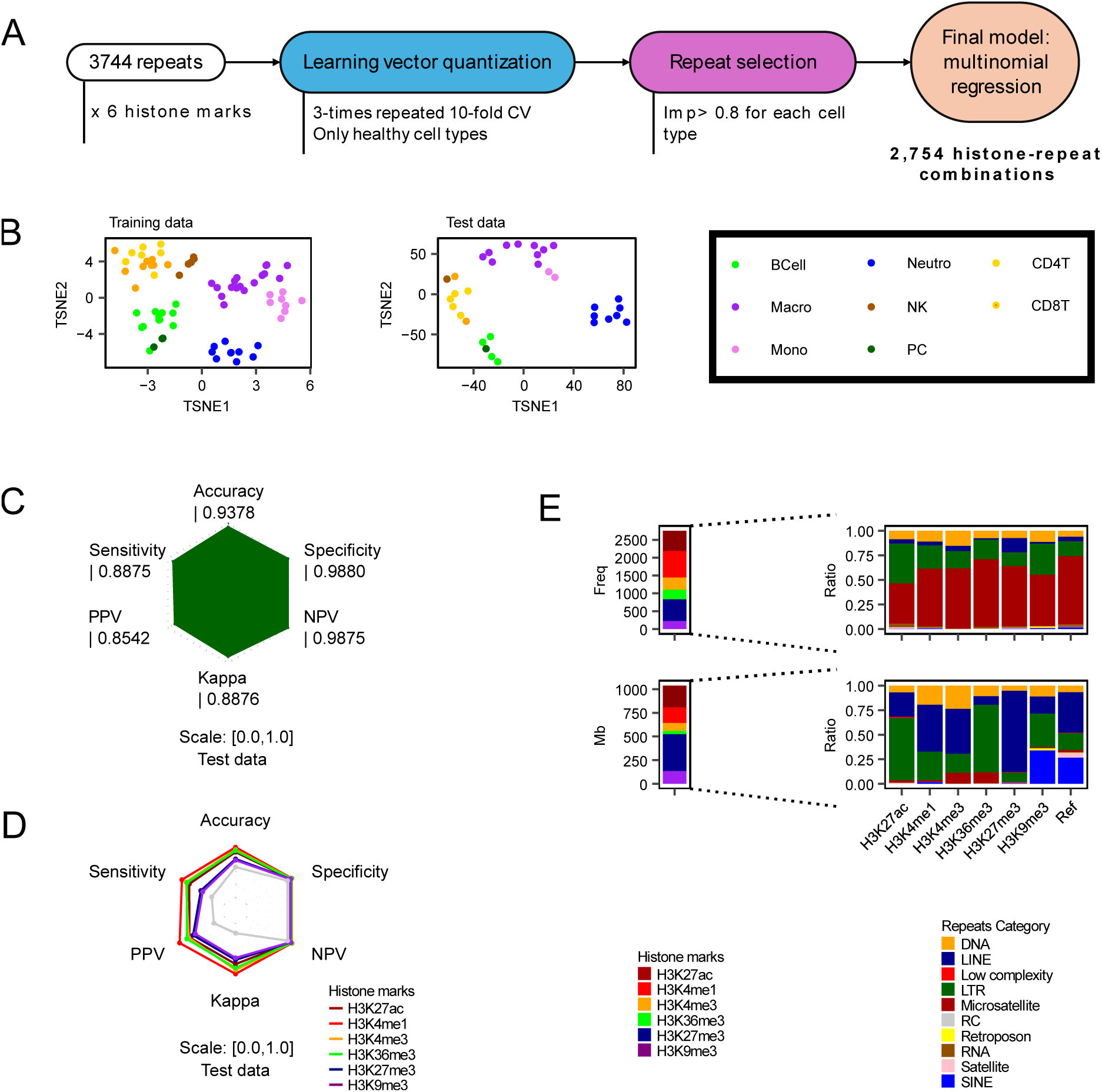
Histone signal–based prediction of cell types in repetitive DNA. (A) Summary of the pipeline used to identify HPTM-RE combinatorial features that predict hematopoietic cell types. (B) t-SNE-based dimensional reduction plots showing sample grouping based on cell type through the use of the 2, 754 HPTM-RE predictors in the training (left) and test data (right). (C) Radar plot showing the performance metrics of the full multinomial regression model in the test data. (D) Radar plot depicting the performance of each HPTM-based multinomial regression model through the test data. (E) Bar plots representing the numbers of HPTM-RE features mapped to each HPTM (top left) and to each RE category separated by HPTM (top right). At the bottom, bar plots showing the genomic coverage of HPTM-RE features mapped to each HPTM (bottom left) and to each RE category, stratified by HPTM (bottom right).

**Supplementary Fig. 5.**
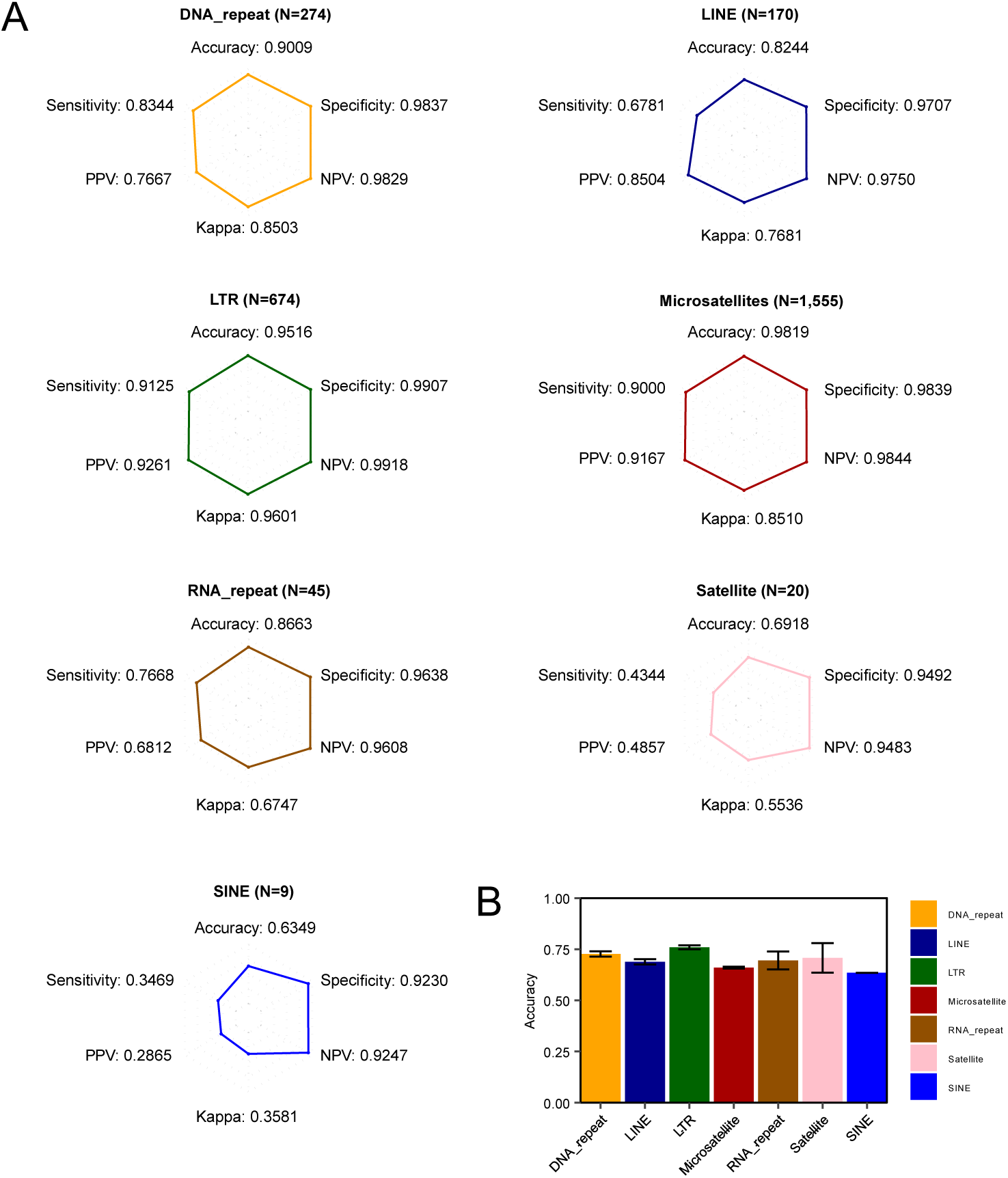
Cell type-specific chromatin information is degenerated across repetitive elements. (A) Radar plots showing the performance metrics for multinomial regression models on test data based on feature sets defined by main RE categories. (B) Bar plots depicting the average test accuracy across all potential multinomial regression models built with 9 predictors per RE category, with standard deviations indicated by error bars.

**Supplementary Fig. 6.**
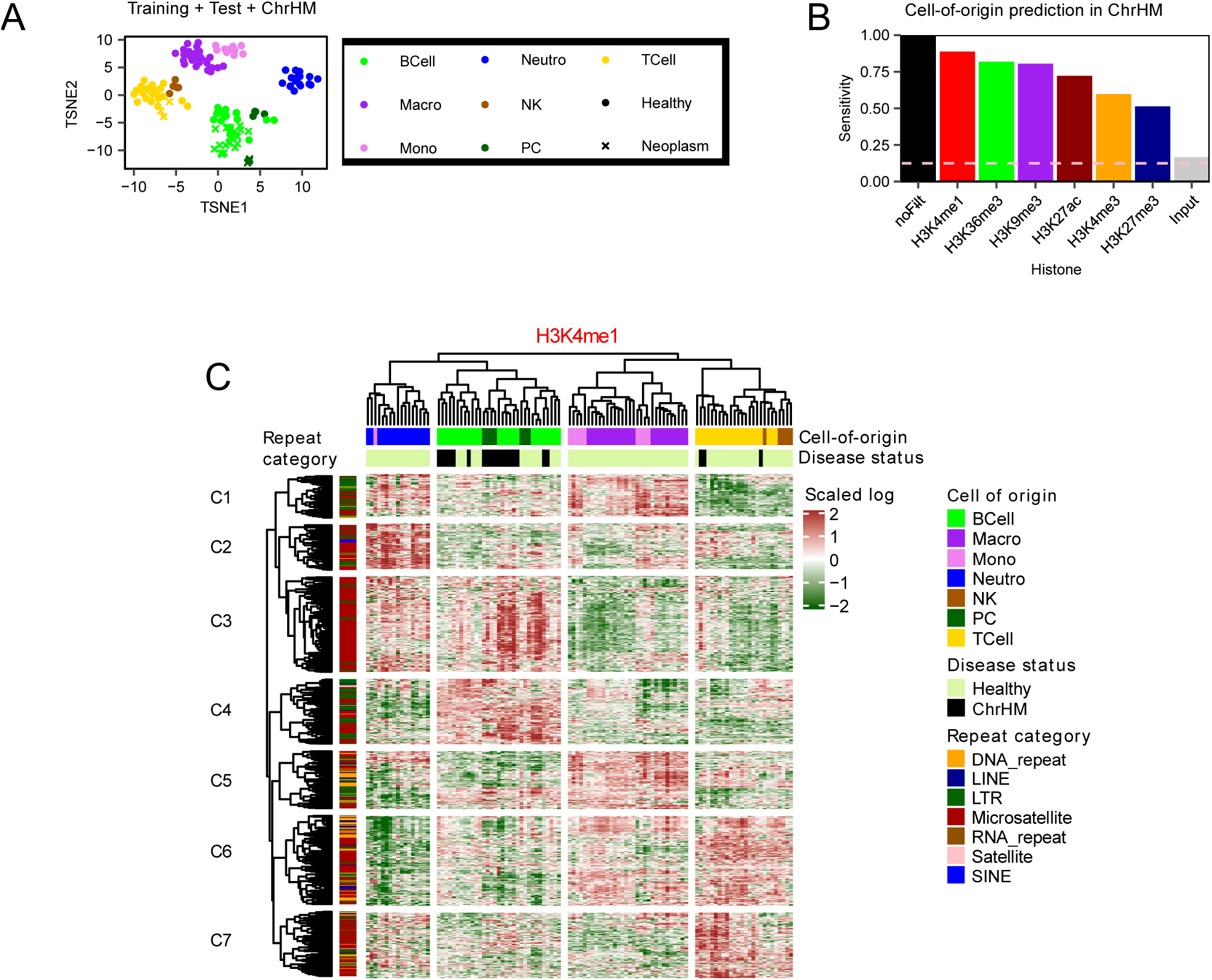
Chronic hematological malignancies preserve repetitive DNA chromatin signatures of their healthy cells of origin. (A) t-SNE-based dimensional reduction plots depicting training, test and leukemic samples clustered by the 2, 754 HPTM-RE cell-type predictive features. (B) Bar plots showing the sensitivity of HPTM-based multinomial regression models in predicting cell of origin in ChrHM, including the negative control (input), the prediction obtained using the full set of HPTM-RE features (noFilt), and the sensitivity expected by random chance (dashed pink line). (C) Heatmap representing scaled, *vst* log-transformed chromatin signals across H3K4me1-RE features, revealing clear groupings of both healthy and leukemic samples according to their cell of origin. RE features are grouped in 7 clusters in an unsupervised manner according to their distinct H3K4me1 signal patterns across samples.

**Supplementary Fig. 7.**
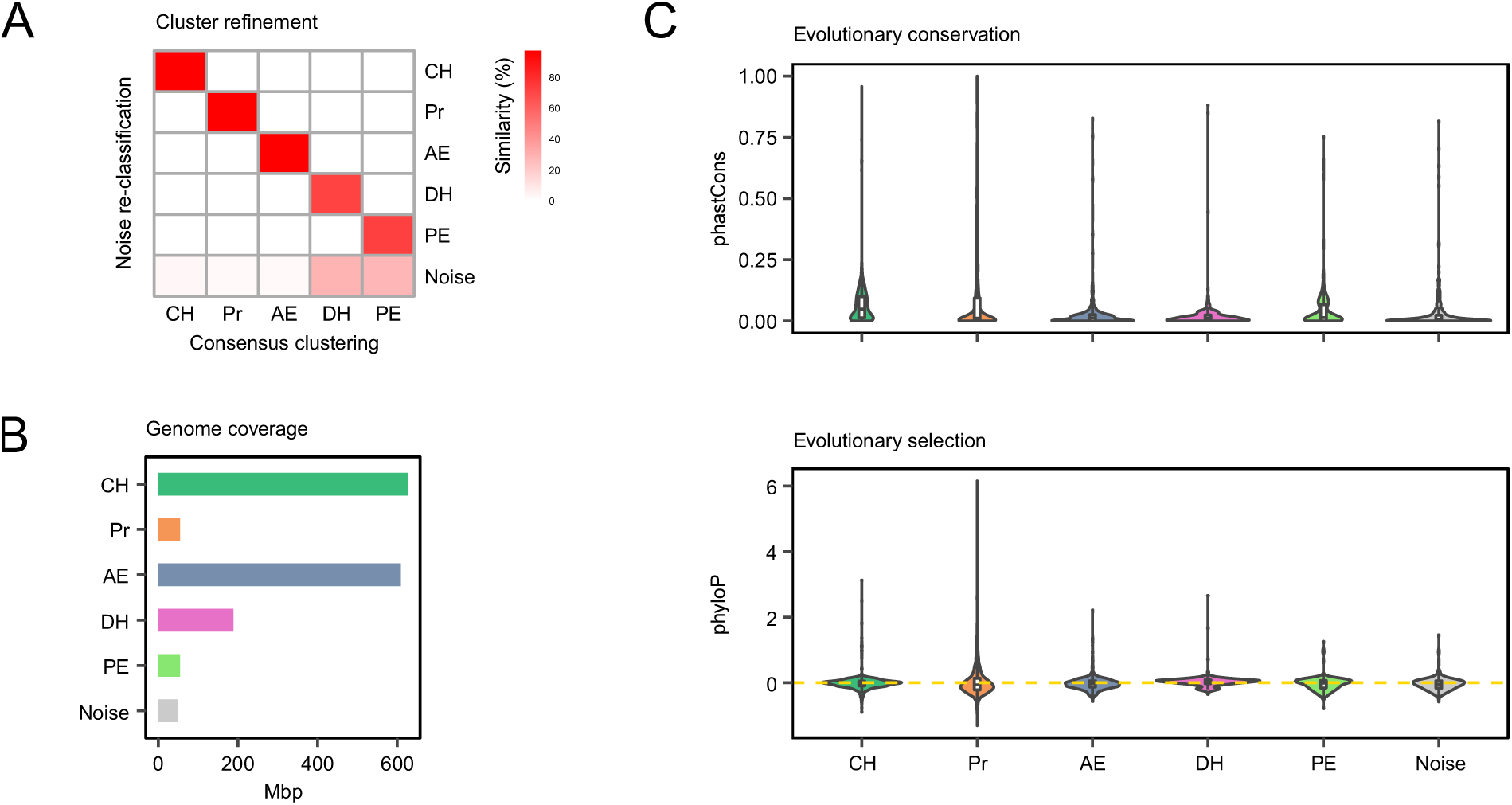
Expanded information on chromatin epitypes. (A) Heatmap showing the percentage of REs within each consensus-clustering epitype that were reassigned to the “Noise” category. (B) Bar plots showing the length of the human genome covered by each RE epitype (Mbp: mega base pair). (C) Violin plots showing the distribution of phastCons (evolutionary conservation) and phyloP (evolutionary selection) scores across RE epitypes.

**Supplementary Fig. 8.**
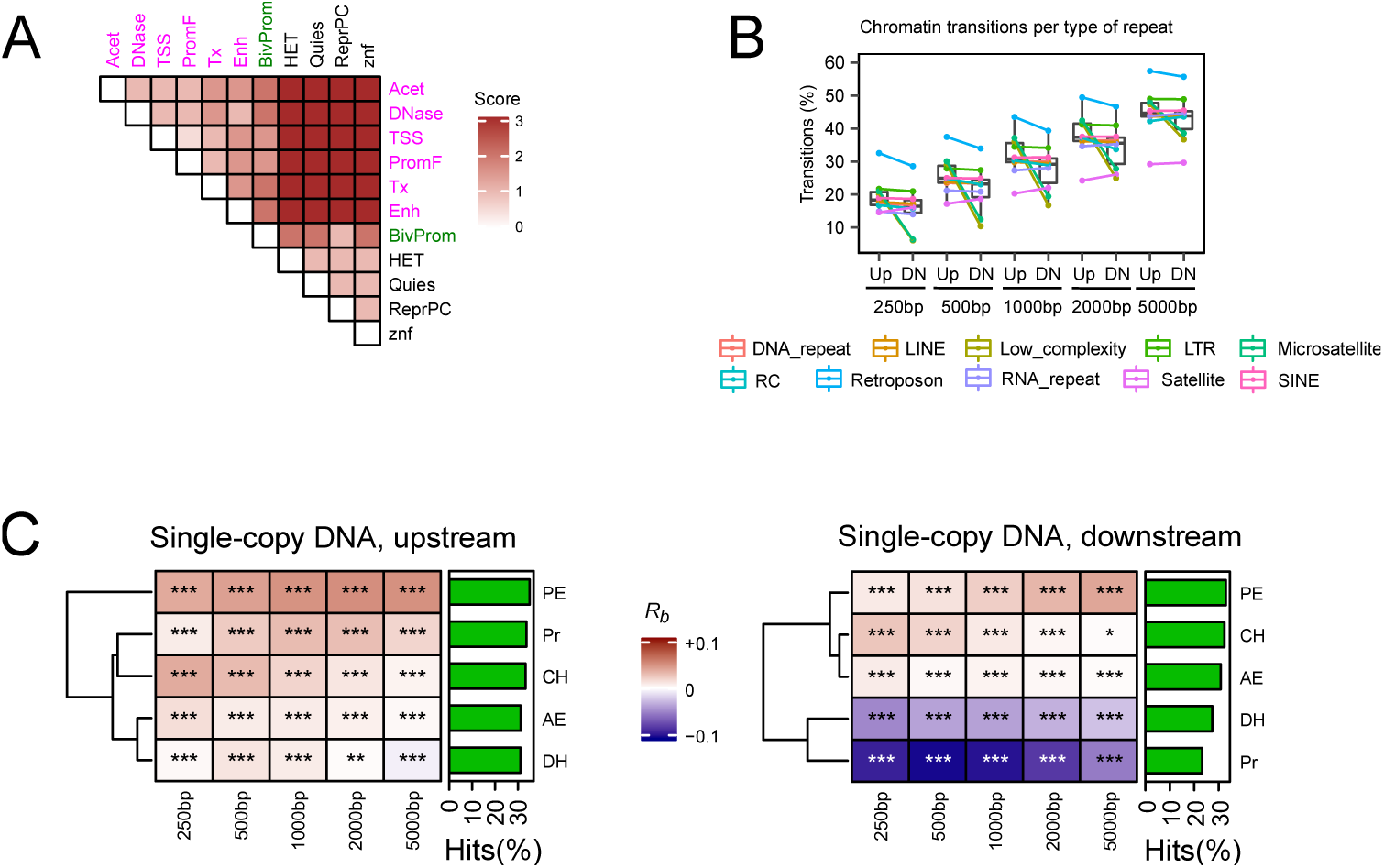
Repetitive DNA chromatin transitions across epitypes and surrounding single-copy regions. (A) Heatmap showing the transition matrix that assigns scores to each RE instance and its surrounding single-copy sequences by comparing their respective chromatin states. (B) Boxplots representing the percentage of REs that undergo any transition per RE category and distance to the RE instance. Lines connect the percentage of transitions for each RE category between upstream and downstream surrounding single-copy sequences. (C) Heatmaps illustrating rank-biserial correlations (*R_b_*) from Wilcoxon rank-sum tests comparing chromatin transition score vectors for each RE epitype against those of all other REs (***FDR *<* 0.001, **FDR *<* 0.01, *FDR *<* 0.05 for two-sided Wilcoxon rank sum tests). Transition scores were computed by contrasting the chromatin state of each RE instance with that of its surrounding single-copy sequences, using the values shown in Supplementary Fig. 8A. On the right side of each heatmap, bar plots indicate the average percentage of REs exhibiting any chromatin state transition across all analyzed intervals (250bp, 500bp, 1000bp, 2000bp, 5000bp).

**Supplementary Fig. 9.**
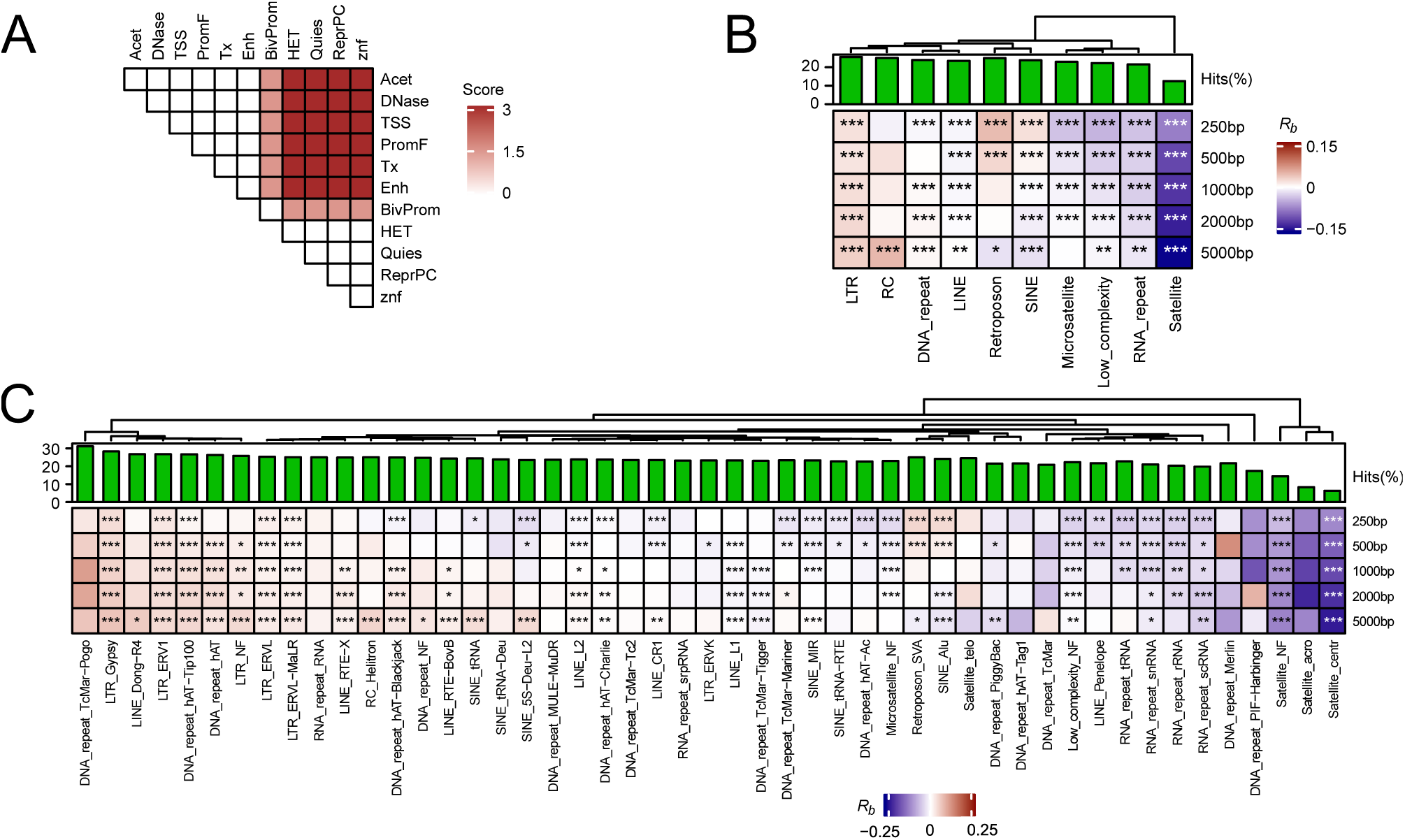
Comprehensive mapping of chromatin boundary sequences in repetitive DNA. (A) Heatmap showing the transition matrix that assigns insulator transition scores to each RE instance by comparing euchromatin and heterochromatin chromatin states between upstream and downstream surrounding single-copy sequences. (B-C) Heatmaps illustrating rank-biserial correlations (*R_b_*) from Wilcoxon rank-sum tests comparing insulator transitions score vectors for each RE category (B) or family (C) against those of all other REs (***FDR *<* 0.001, **FDR *<* 0.01, *FDR *<* 0.05 for two-sided Wilcoxon rank sum tests). Insulator transition scores were computed by contrasting per RE instance the chromatin state of upstream and downstream surrounding single-copy sequences, using the values shown in Supplementary Fig. 9A. On the right side of each heatmap, bar plots indicate the average percentage of REs exhibiting any insulator transition across all analyzed intervals (250bp, 500bp, 1000bp, 2000bp, 5000bp).

**Supplementary Fig. 10.**
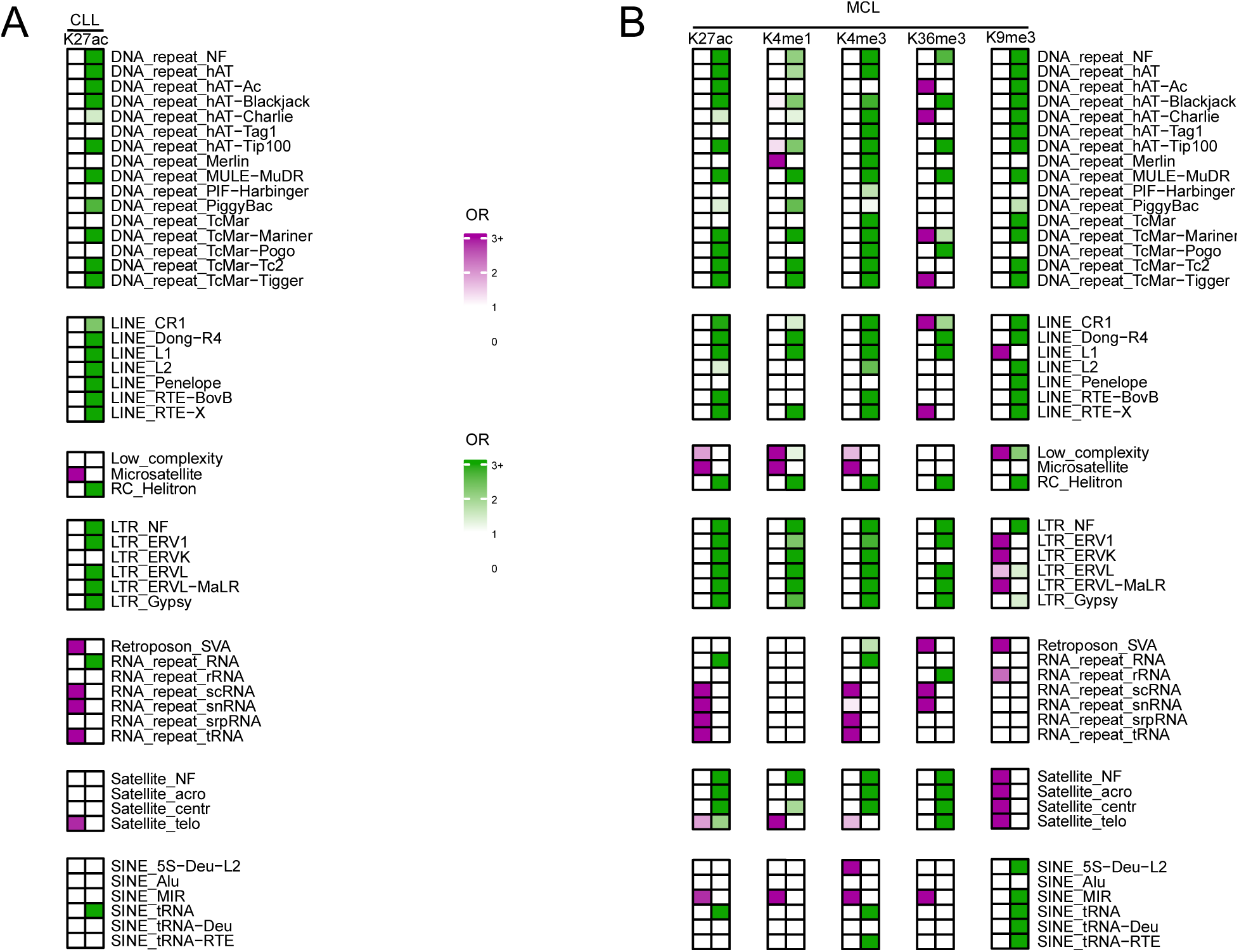
Dynamic remodeling of histone signals across repetitive element families in B-cell chronic malignancies. Heatmaps representing the enrichment (odds ratios: OR) of significantly upregulated and downregulated RE subfamilies (FDR<0.05) across RepeatMasker-annotated families and HPTMs, in CLL (A) and MCL (B).

**Supplementary Fig. 11.**
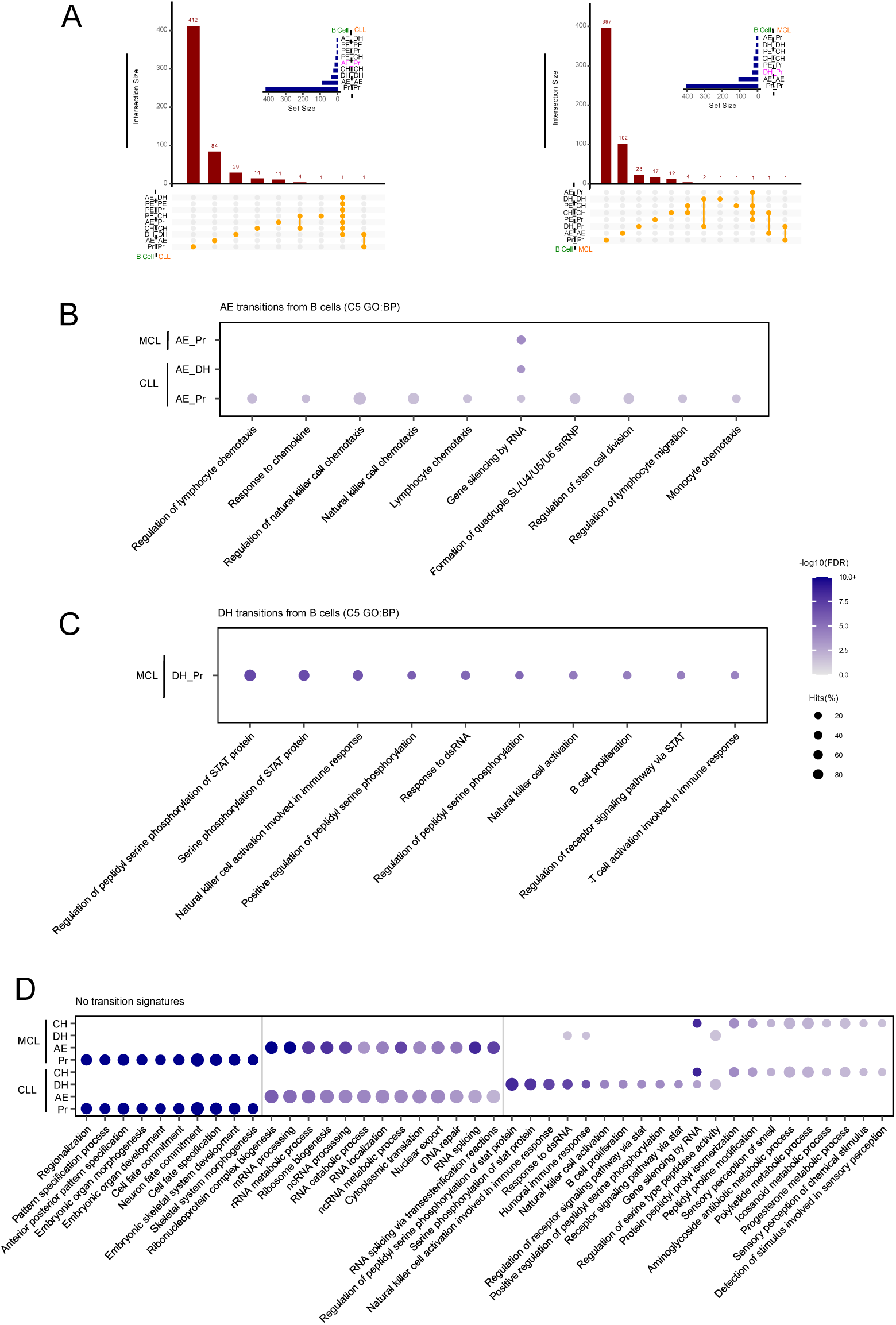
The functional landscape of genes enriched in repeats undergoing chromatin epitype switches. (A) Upset plots showing the number of significantly enriched ontologies (FDR<0.05, C5 GO BP) associated with the pairwise combinations of epitypes observed between B cells and CLL (left) or MCL (right). Most important transitions are colored in violet. (B-D) Bubble plots depicting the top 10 most significant gene ontologies associated with: (B) AE transitions in CLL and MCL, (C) DH to Pr transitions occurring in MCL; and (D) no epitype transitions between B cells and ChrHM (D).

**Supplementary Fig. 12.**
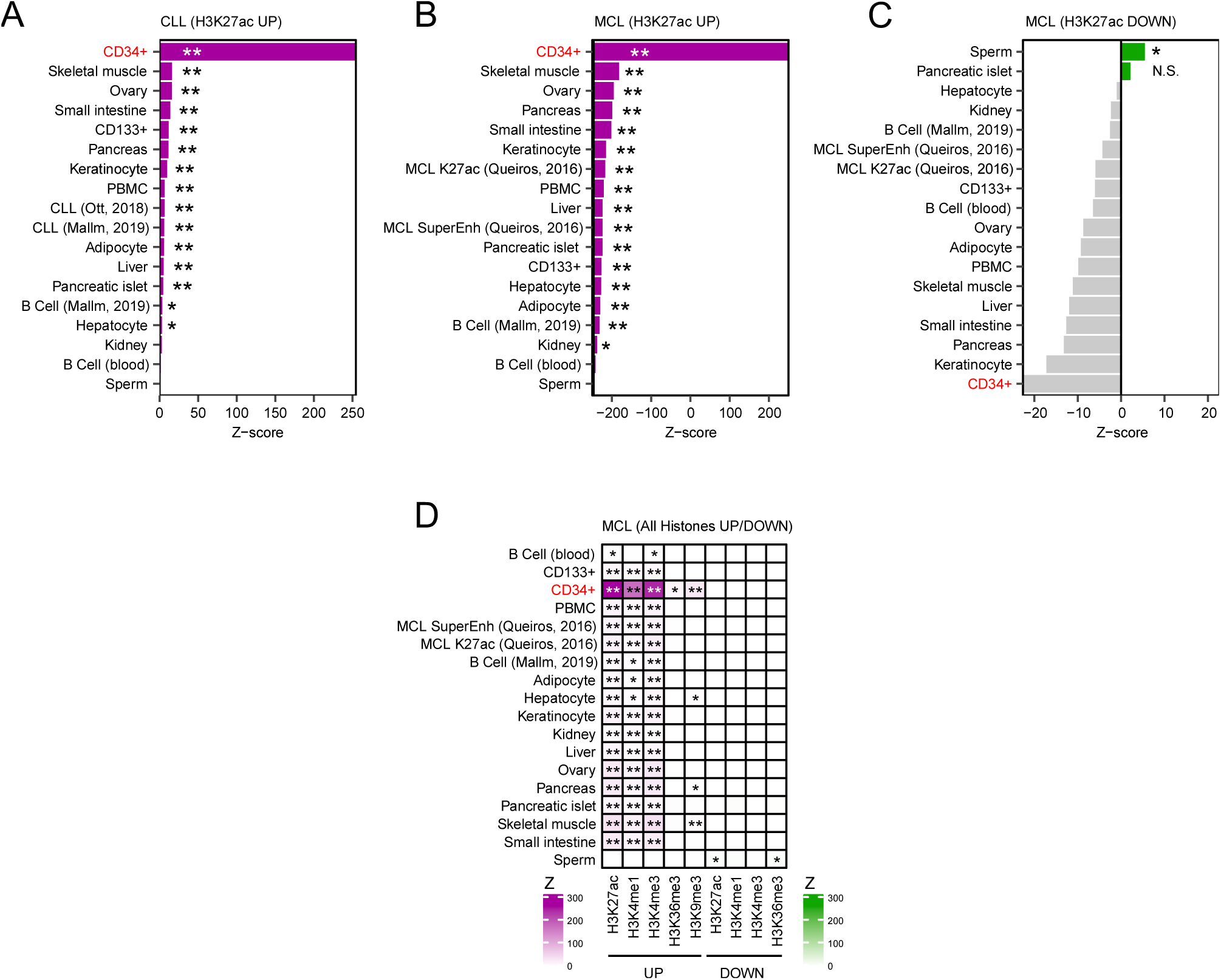
Characterization of enhancer regulatory sequences enriched in altered repetitive elements. (A-C) Bar plots showing enrichment scores (where Z is the z-score obtained for one-sided permutation *regioneR* tests) derived from overlaps between distinct human enhancer regions and (A) upregulated H3K27ac REs in CLL (FDR<0.05, log_2_(FC)*>*1); (B) upregulated H3K27ac REs in MCL (FDR<0.05, log_2_(FC)*>*1); and (C) downregulated H3K27ac REs in MCL (FDR<0.05, log_2_(FC)*<* –1). Positive Z-scores are colored in purple for upregulated REs and in green for downregulation REs; whereas negative Z-scores are shown in grey. (D) Heatmap of enrichment scores (where Z is the z-score obtained for one-sided permutation *regioneR* tests) depicting overlaps between distinct enhancer regions and significantly altered RE subfamilies per HPTM (FDR<0.05, |log_2_(FC)|*>*1), stratified by direction of change (purple: upregulated REs; green: downregulated REs; negative Z-scores are set to white).

**Supplementary Fig. 13.**
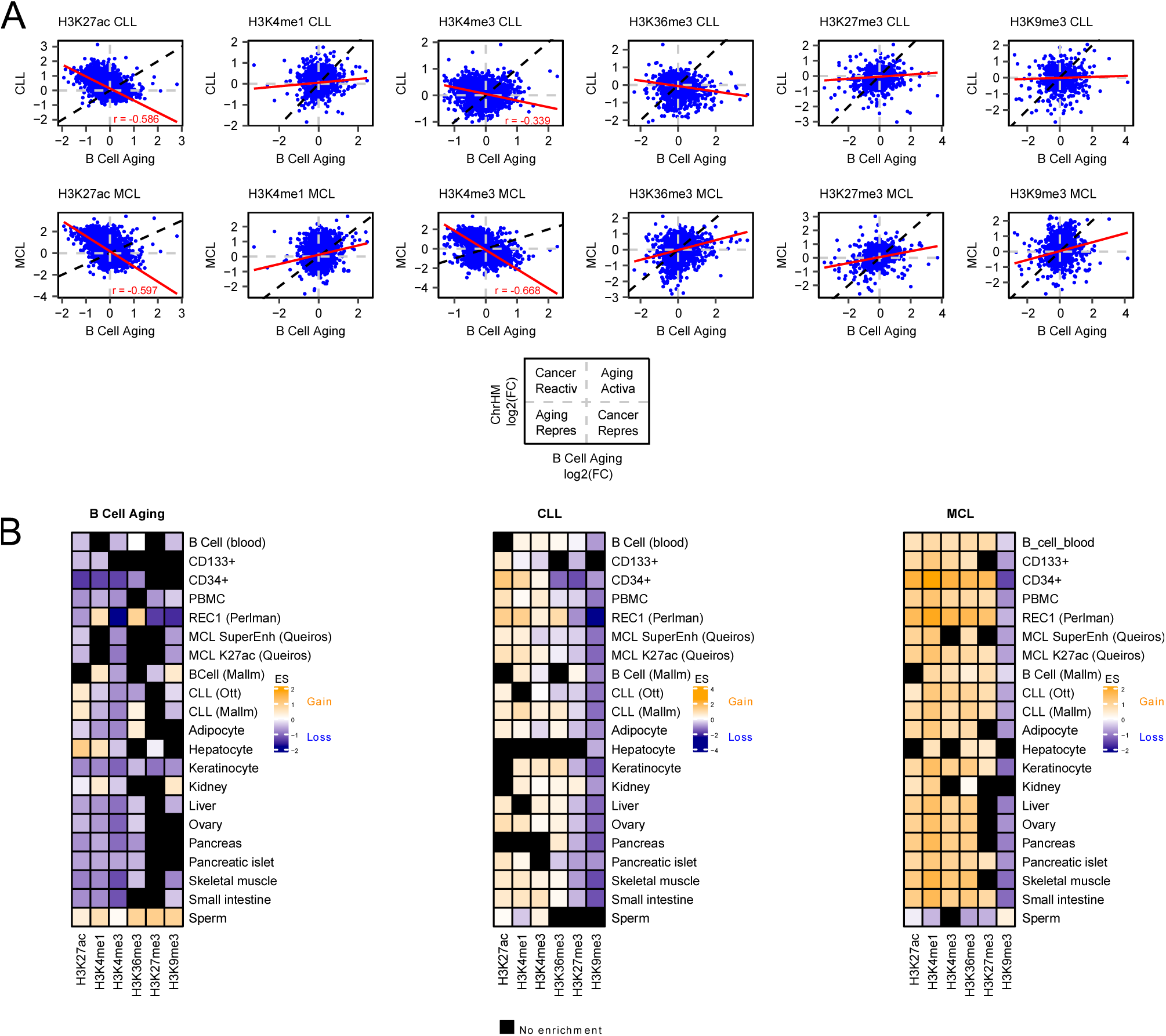
Cancer reverses age-associated alterations in active HPTMs within the B-cell compartment. (A) Scatter plots showing the Pearson correlation between the log_2_ fold changes of all REs (3,744) in B cell aging and malignant transformation (CLL [top] or MCL [bottom]) per HPTM. Perfect linear positive correlations are represented using black dashed lines. (B) Heatmap of enrichment scores (sign(Z_diff_) × log_10_(|Z_diff_|+1), where Z_diff_ = Z_up_ − Z_down_) depicting gains (>0) or losses (<0) of HPTM signals in REs overlapping enhancer regions during B cell aging (left) and malignant CLL (middle) or MCL (right) transformations. Z is the z-score obtained for one-sided permutation regioneR tests with the 50 RE subfamilies with the lowest *p*-values, per HPTM and direction of change. If Z_up_<0 and Z_down_>0, the selected REs are less enriched than expected within enhancer regions, and color is set to black.

## Supplementary table legends

**Supplementary Table 1.** List of ChIP-seq experiments analyzed in this study, annotated by HPTM and the corresponding purified hematopoietic cell type or chronic hematological malignancy.

**Supplementary Table 2.** Identification of REs with a significantly higher signal in unprecipitated chromatin samples (*replist*), using the *DESeq2* suite.

**Supplementary Table 3.** Classification of RE subfamilies according to chromatin epitypes.

**Supplementary Table 4.** Differential HPTM signal alterations in REs during CLL malignant transformation from B cells, as determined by *DESeq2* analysis.

**Supplementary Table 5.** Differential HPTM signal alterations in REs during MCL malignant transformation from B cells, as determined by *DESeq2* analysis.

**Supplementary Table 6.** Differential HPTM signal alterations in REs during B-cell aging, as determined by *DESeq2* analysis.

